# Replay of procedural experience is independent of the hippocampus

**DOI:** 10.1101/2024.06.05.597547

**Authors:** Emmett J. Thompson, Lars Rollik, Benjamin Waked, Georgina Mills, Jasvin Kaur, Ben Geva, Rodrigo Carrasco-Davis, Tom George, Clementine Domine, William Dorrell, Marcus Stephenson-Jones

## Abstract

Sleep is critical for consolidating all forms of memory^1-3^, from episodic experience to the development of motor skills^4-6^. A core feature of the consolidation process is offline replay of neuronal firing patterns that occur during experience^7,8^. This replay is thought to originate in the hippocampus and trigger the reactivation of ensembles of cortical and subcortical neurons^1,3,9-18^. However, non-declarative memories do not require the hippocampus for learning or for sleep-dependent consolidation^19-26^ meaning what drives their consolidation is unknown. Here we show, using an unsupervised method, that replay occurs in the dorsal striatum of mice during offline consolidation of a non-declarative, procedural, memory and that this replay is generated independently of the hippocampus. Replay occurred at both real-world and time-compressed speeds and was also prioritised both at the level of the individual neurons and the type of neural sequence. Complete bilateral lesions of the hippocampus had no effect on any feature of this replay. Our results demonstrate that procedural replay during consolidation of a non-declarative memory is independent of the hippocampus. These results support the view that replay drives active consolidation of all types of memory during sleep but challenges the idea that the hippocampus is the source of this replay.

---

The hippocampus is believed to be critical for the active consolidation of all types of memory as it is thought to trigger reactivation of cortical memory traces that, when strengthened over time can be activated independently of the hippocampus^1,27,28^. However, the hippocampus is not required for the formation of many forms of non-declarative memory^19-26^. Indeed, lesions of the hippocampus may even facilitate procedural or associative memory formation^20,24,26,29^. This raises the questions as to how memories such as those for procedural skills are consolidated during sleep and whether there is a source of replay independent of the hippocampus.

## Development of a multi-step procedural memory task

To investigate the mechanisms that support offline consolidation of procedural memory we developed a multi-step procedural memory task for mice. Mice were rewarded for nose-poking in the correct sequence of five ports (Fig.1a and Extended Data Fig.1a,b and Supplementary Video 1). Mice learnt to produce the sequence from memory (Fig.1b) and once experts, completed the full task with highly stereotyped timing and accuracy (Fig.1c,d and Extended Data Fig.1c). Trial-to-trial movement variability decreased across learning, (Extended Data Fig.1d) consistent with the notion that animals had acquired a stereotyped procedural skill. Motor skill learning and execution is thought to depend on the dorsolateral striatum (DLS)^30,31^. To confirm that the DLS was important for learning and executing our multi-step task we lesioned this structure prior to training using a viral caspase strategy (Fig.1e,f and Extended Data Fig.2a-c). Compared to control mice, DLS lesioned animals showed impaired task learning (Fig.1g,h and Extended Data Fig.3a). The learning curves between groups diverged at a training level where visual guidance was removed, indicating that the DLS is needed for learning to perform the sequence from memory but not in a light guided manner. This effect was specific to the DLS as mice with lesions to the adjacent dorsomedial striatum (DMS) did not show learning impairments (Extended Data Fig.3b-e). Post-learning DLS lesions or inactivation impaired task performance and increased movement variability (Fig.1i-k and Extended Data Fig.3f,g,i), without affecting individual motor elements or port relevance recognition (Extended Data Fig.3h). Taken together, these results show that the DLS is required for learning and executing the motor sequence from memory.

**Figure 1:**
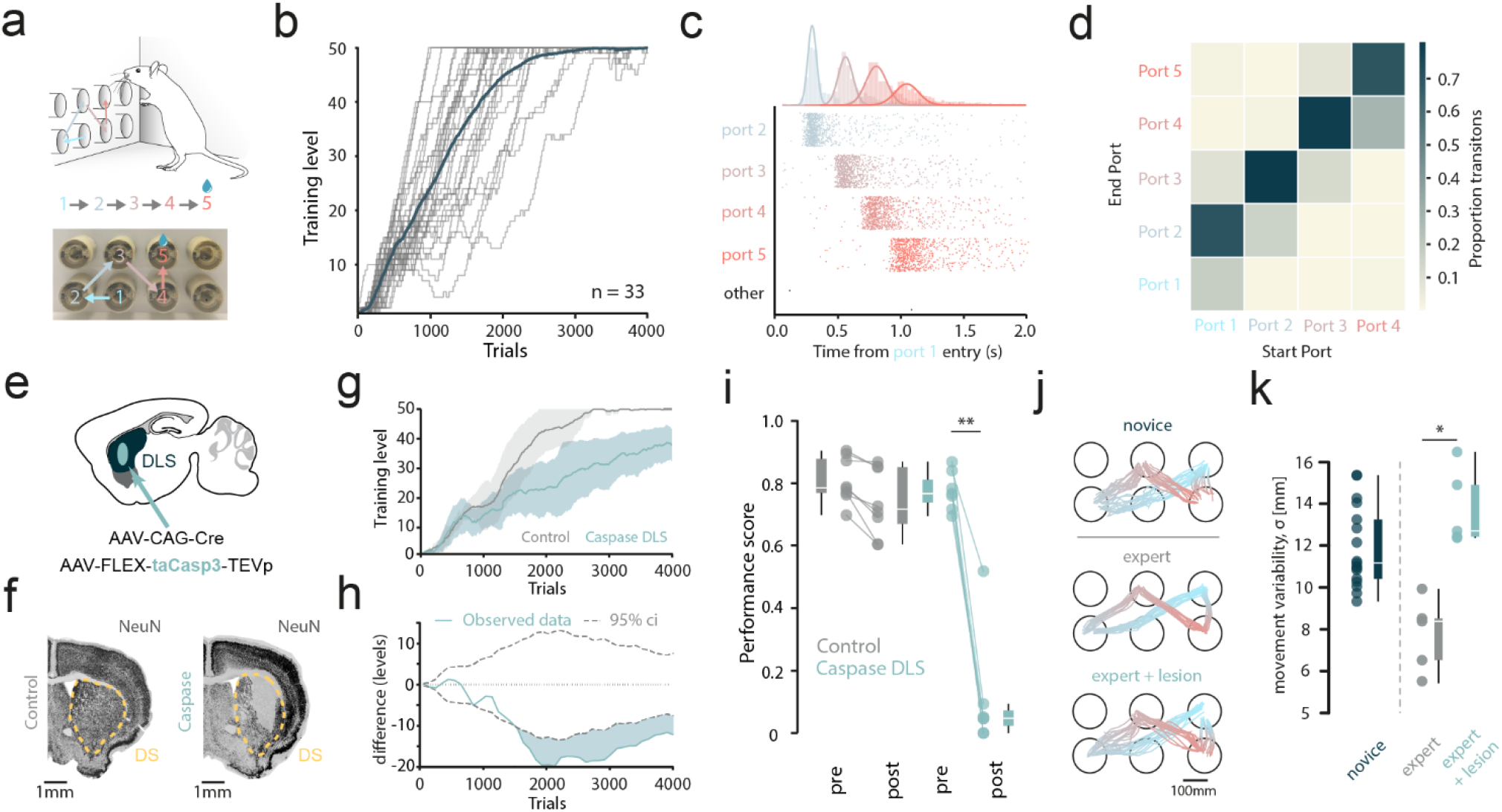
Learning and execution of a novel sequence learning task is dependent on DLS. **a**. Top: schematic of the task. Bottom: Photograph of the port wall with task sequence overlaid. **b**. Training level progression curves (grey) for multiple animals with mean learning curve overlaid (dark green) (n = 33 animals, mean trials to level 50 = 1942 +/-111; SEM). **c**. Example poke times from an expert animal. Port-poke-in times are shown (points) for each port relative to the start of trial initiation. All pokes in task irrelevant ports are labelled ‘other’. **d**. Transition heatmap showing mean port to port transition proportions for multiple trained animals (n = 33). Each transition in the task sequence is represented by start (x-axis) and end (y-axis) ports. **e**. Schematic of the experimental approach for bilateral lesion of the DLS. **f**. Histology; coronal sections showing neurons in one example hemisphere of the striatum (yellow outline) for lesion and control mice. **g**. Learning curve (training level vs trials) for control and lesion animal groups (shaded area denotes standard deviation, lesion n = 7 mice, control, n = 6 mice). **h**. Differences in performance between the groups. Dotted lines indicate the 95% confidence interval for the shuffled data (see methods). **i**. Average task performance scores (see methods) for the 3 sessions pre and post injection surgery (Kruskal-Wallis; p = 0.0006. For displayed stars, Post-hoc Dunn test; p = 0.002. lesion, n = 7 mice; control, n = 8 mice). **j**. Movement tracking (animal head) for 4 subsequence movement vectors for an example novice, expert and lesioned expert mouse. **k**. Average movement variability (standard deviation from mean trajectory) across the four subsequence trajectories for novice mice (n = 8 mice, n = 18 sessions) and expert mice before and after lesion to DLS (paired t-test; p = 0.0085, n = 5 mice).

### NMDA dependent plasticity supports offline procedural memory consolidation in the DLS

To determine if the DLS is involved in offline consolidation, we infused the NMDA receptor antagonist 2-amino-5-phosphonopentanoate (AP5) into the DLS to disrupt offline striatal activity and plasticity^32-34^ . This was done immediately after the days training, before placing mice back into their home cage to sleep ^32-34^ (Fig.2a,b, Extended Data Fig.4a). Testing the next day, 24 hours after infusion, across all infusion sessions there was a significant effect of AP5 infusion on performance (Fig.2c). In early training, mice in the AP5 cohort were significantly worse at the task the following day (Fig.2d), on average they dropped to a training level they had been at the day before the infusion session (Extended Data Fig.4b). This suggests that offline NMDA-dependent plasticity in the DLS is critical for consolidating the previous day’s performance gains. To determine if offline activity and plasticity was also critical for maintaining procedural memory, we infused saline or AP5 for 4 consecutive days in the DLS post-training (Extended Data Fig.4c). Infusion of AP5 for 4 consecutive days led to a significant drop in performance (Extended Data Fig.4d). Together these data suggest that offline processes in the dorsolateral striatum are critical for both learning and maintaining procedural memory of a stereotyped action sequence.

**Figure 2:**
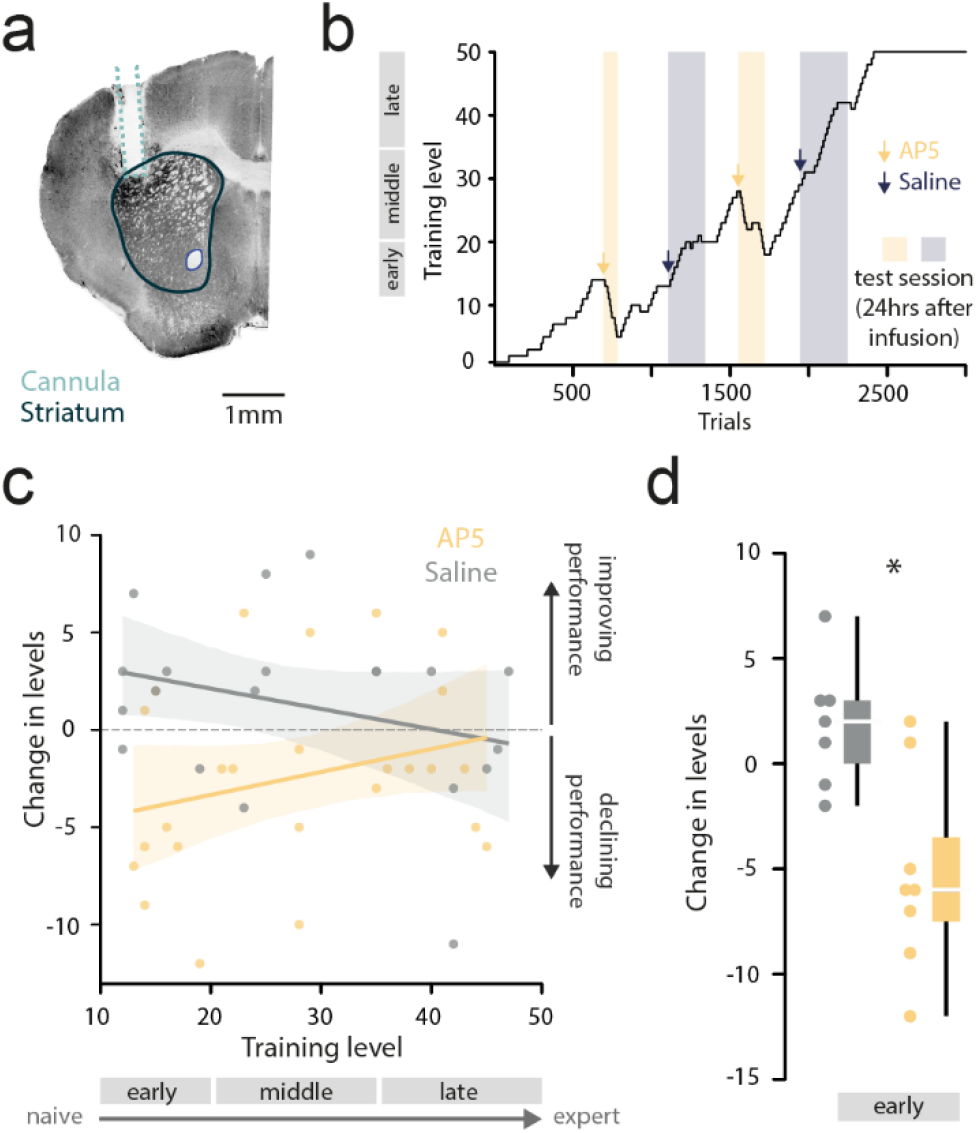
Blocking offline plasticity and activity in the DLS impairs learning of procedural memory. **a**. Histology showing an example hemisphere of a coronal slice after bilateral cannula implantation. Cannula tract (light blue) and striatum (dark blue) are outlined. **b**. Example animal learning curve showing AP5 and saline infusions (arrows) and test sessions 24 hours after infusion (shaded). Infusions were performed the day before the test session, immedialty after training that day (see methods) **c**. Training level changes against the start level for all post infusion test sessions (saline and AP5) for all animals (n = 8). Lines show linear regression fit for each dataset with confidence interval (shaded region) (p = 0.033, Ordinary Least Squares regression). **d**. Training level change for early learning infusion experiments (p = 0.00448, independent t-test).

### An unsupervised approach for replay detection

If an offline process supports procedural memory formation in the DLS, what could the underlying mechanism be? For episodic memories, replay of previously observed neural activity is thought to be the way in which memory is consolidated offline^7,8^. To search for similar neural reactivations in the DLS we implanted neuropixel probes through this region and recorded neural activity during task execution and post task sleep (Fig.3a). Typically replay has been identified using template-based approaches such as template matching or Bayesian decoding^35-38^. These approaches often carry assumptions, such as that neural sequences progress in a linear fashion, in a constant direction at a constant speed. These assumptions have been shown to limit the types of replay that have been detected in the hippocampus^39^. In the hippocampus, these methods are also usually applied during candidate epochs defined by the presence of sharp-wave ripple (SWR) oscillations which are thought to be a marker for hippocampal replay^40^. As we did not know of a heuristic biomarker for replay in the striatum and did not want to a priori limit the types of reactivations that we could detect we adapted an unsupervised point process model called PP-Seq^41^.

**Figure 3:**
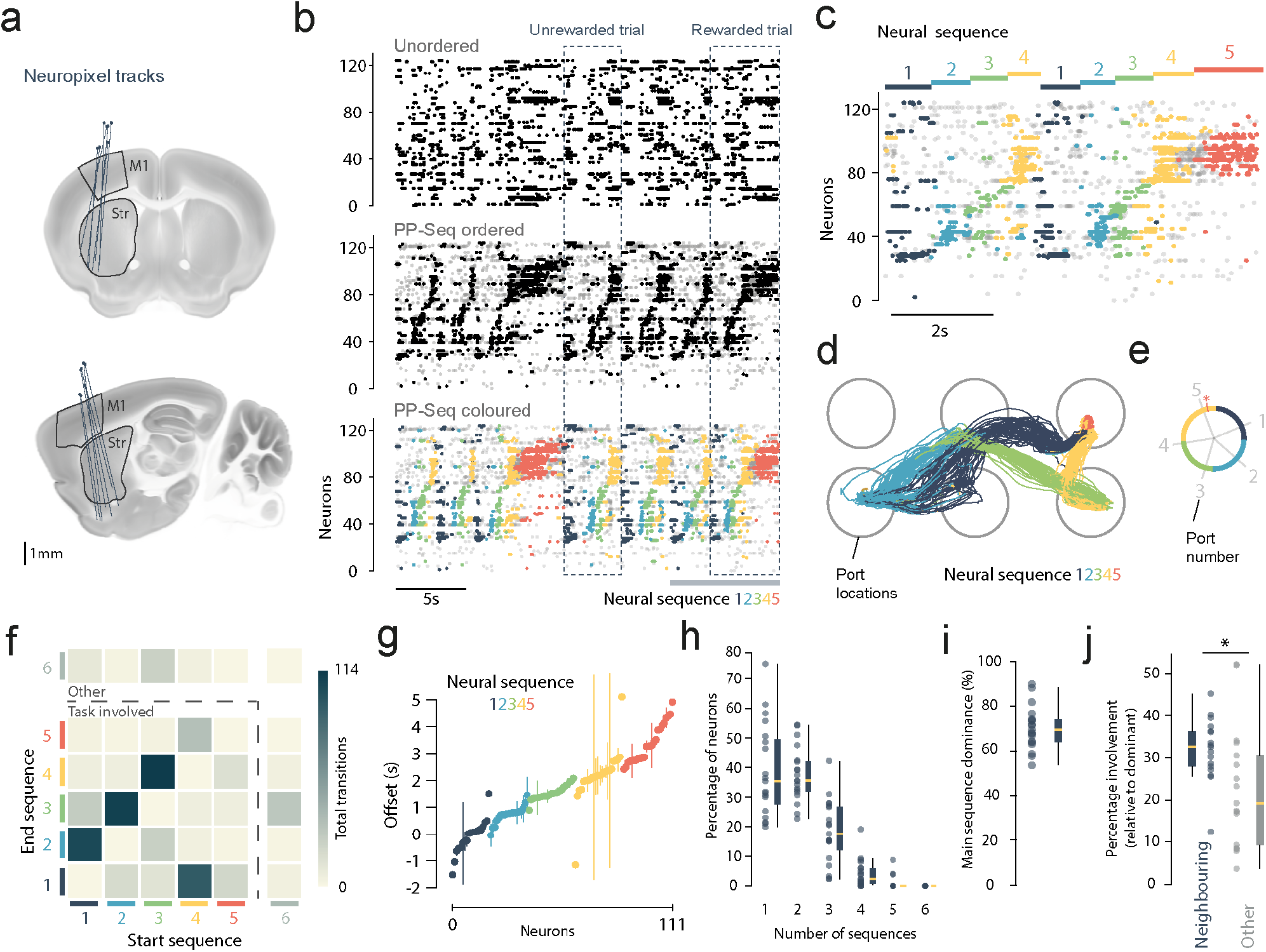
Unsupervised labelling of latent neural sequences in DLS during task practice. **a**. Schematic showing implanted neuropixel probe locations projected onto standard Allen atlas coronal (top) and sagittal (bottom). **b**. Top: unordered example spike raster showing spikes for neurons in striatum recorded during task execution. Middle: the same spike raster but neurons ordered lexicographically based on PP-Seq identified neural structure. Bottom: Same as above but spikes coloured by which latent PP-Seq sequence they were attributed to. A rewarded and an unrewarded trial are shown (dashed boxes). **c**. Example PP-Seq ordered spike raster (as in panel b, grey bar bottom right) zoom in, showing two task trials with task relevant sequences labelled. **d**. Example movement tracking (animal head) from 400s of task engagement, coloured by the current dominant PP-Seq sequence. **e**. Circularised task space coloured by the dominant PP-Seq labelled neural sequence. Hidden (non-dominant on average) task relevant sequence is represented by a star. Grey lines indicate respective task port locations across standardised space. **f**. Transition histogram showing numbers of sequence-sequence transitions during the example epoch. Task associated and non-task associated sequences are separated by the dotted line. **g**. Mean spike times during instances of each neuron’s dominant PP-Seq sequence for the example animal (Error bars show standard deviation over sequence instances). Neurons are ordered by offset from the earliest neuron in each sequence and sequences ordered by mean distance between sequence midpoints. **h**. Number of sequences neurons appeared in. For each analysed recording session, the percentage of neurons that contribute (appearing at least 10% of the time) to different numbers of sequences is shown. **i**. Mean relative percentage per session of spikes that were present in their most common (dominant) sequence. **j**. Mean neuron occurrences between neighbouring and distal sequences for neurons in each analysed recording session. Mean proportions are calculated relative to the dominant sequence proportion. Sequence order was most observed task order and neighbouring sequences were defined by those adjacent to the dominant (p = 0.027, paired t-test, n = 19 sessions, n = 7 mice, yellow markers indicate median)

PP-Seq aims to attribute individual neuronal spikes to a latent cause, in this case instances of particular neural sequences. Via an iterative collapsed Gibbs sampling procedure, the model fits free parameters to determine the number of latent events (neural sequences) and then attributes spikes to these sequences based on features of activity (Extended Data Fig.5a). In our raw neuropixel data there were no visible sequential patterns; however, clear sequences were revealed by sorting the neurons lexicographically according to the sequence type and the temporal offset parameter inferred by PP-Seq (Fig.3b,c). Using a hyperparameter search we found that our data was best described as containing six types of neural sequence (Extended Data Fig.6a-c). To determine if these neural sequences were related to aspects of our multi-step task, we aligned these neural sequences with video tracking data and coloured the tracking by the dominant sequence type. Strikingly, despite the model having no access to behavioural information, different types of neural sequences aligned to distinct phases of the behavioural sequence (Fig.3d-f and Supplementary video 2). In every mouse recorded, the behavioural sequence was accompanied by multiple neural sequences that aligned to distinct phases of the task (Extended Data Fig.5b). Not all neural sequences were aligned to distinct movement between the ports, in some animals specific neural sequences correlated with reward consumption or aligned tightly to other behavioural sequences that the mice performed during the recording session such as grooming (Extended Data Fig.5c and Supplementary video 3). The neurons within a sequence type tended to spike at precise timing during the sequence and were also relatively exclusive, mostly spiking in one neural sequence type (Fig.3g-i). When neurons did contribute spikes to multiple types of neural sequence the spikes were most likely to occur in the preceding or following neural sequence (Fig.3j). Taken together, applying PP-Seq revealed that there were multiple neural sequences in the striatum that occurred at distinct phases of the behavioural sequence. This suggests that an aspect of procedural learning may be in learning to chain together these neural sequences that are correlated with individual portions of the behaviour.

### Procedural replay in the DLS

To determine if the neural sequences that we observed in the awake data were reactivated offline during sleep we applied our awake trained PP-Seq model to post task activity recorded during sleep (Fig.4a) Neural sequences that were observed in the awake data were reactivated during sleep (Fig.4b,c) but were not present in shuffled sleep data (neuron IDs permuted, Extended Data Fig.7a). To benchmark our unsupervised method for replay detection we compared it with two Bayesian decoding methods, a classic linear decoding method^36,42^ and a state-of-the-art combined state-space and nonlinear decoding method^39^. On synthetic data, designed to create a ground truth dataset, PP-Seq was both more accurate and more sensitive at detecting replay than either of the decoding methods (Extended Data Fig 7b-e). PP-Seq was also more robust at finding replay when noise, jitter and temporal distortions were applied to the data (Extended Data Fig 7f-i). In our actual recording data, the majority of PP-Seq identified replay events were also identified by the non-linear decoding method, validating that multiple methods identify replay of our task related activity in the striatum during sleep (Extended Data Fig.8a-g). Neural firing during replay followed a sequential ordering at the level of the single neurons (Fig.4d and Extended Data Fig.8h) and this ordering largely matched that observed for the same neurons during awake sequences (Fig.4e and Extended Data Fig.8i). At a macro scale, neural sequences were often replayed individually but also occurred in combination (Fig.4f,g). When combinations of sequential neural sequences were replayed, they were more likely than chance to occur in the order that they appeared in the awake data as mice performed the behavioural sequence (figure 4h). This shows that both the sequential firing within a neural sequence and the compositional order between neural sequences was maintained during replay.

**Figure 4:**
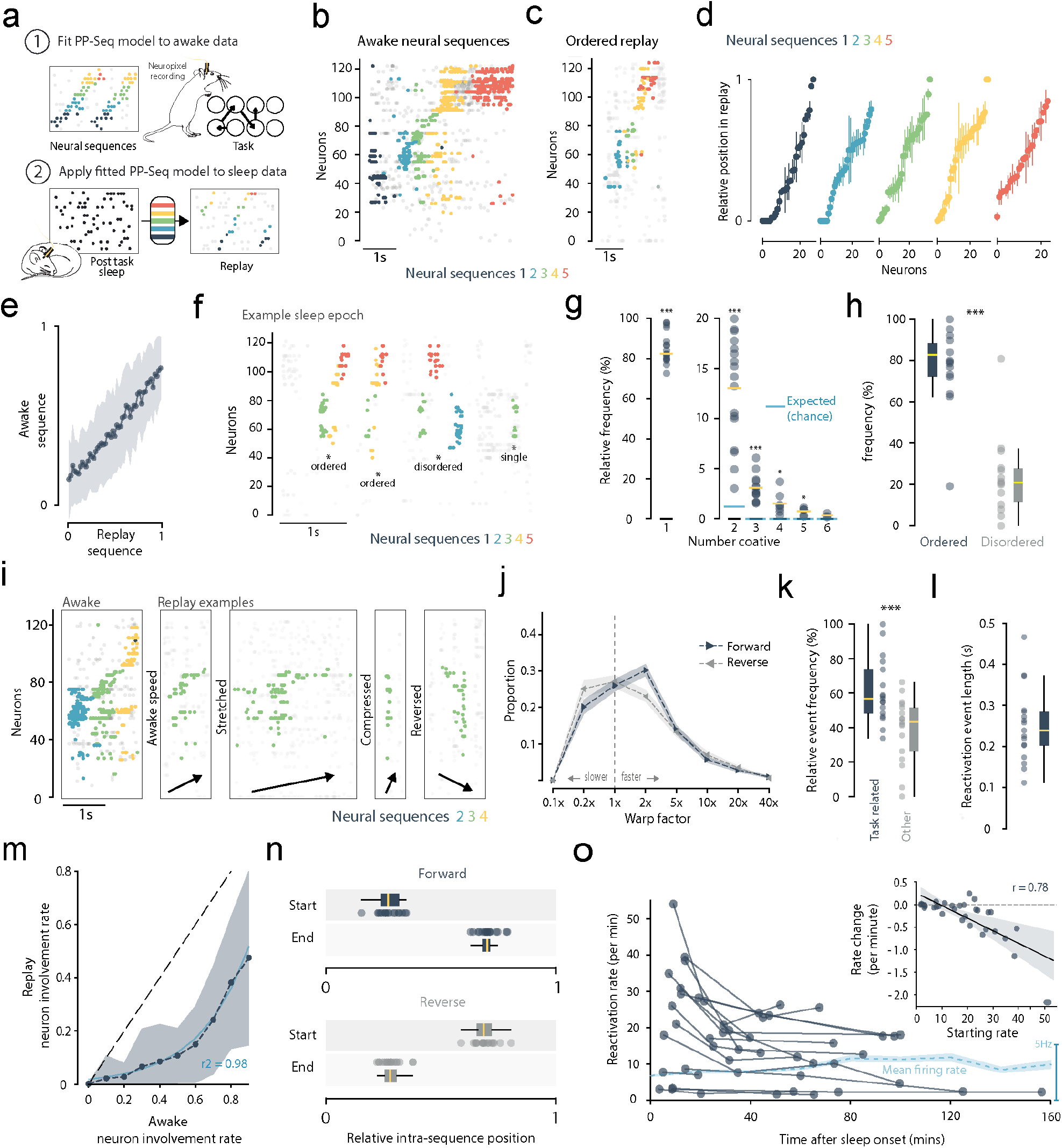
Replay of procedural neural activity in the striatum during sleep. **a**. Schematic of replay detection protocol. 1: PP-Seq model is trained on and detects repeating neural sequences within task related activity. 2: Fitted model is applied to sleep recording to identify recapitulated task related neural sequences. **b**. PP-Seq labelled sequences for example task related awake spikes. **c**. Example replay (same neurons as in b) of concatenated sequences that are ordered with respect to task related activity. **d**. Mean relative position of spikes, for neurons in each replay sequence observed during the example recording (error bars, SEM). **e**. Mean relative positions in awake sequences vs. replay sequences for all analysed neurons across every session (OLS regression, r = 0.64, p < 0.001). **f**. Example sleep period with PP-Seq labelled replay sequences **g**. Frequency of single (isolated) and concatenated events for each recording session. (difference from chance, one sample t-tests, left to right; p < 0.001, p < 0.001, p < 0.001, p = 0.0117, p = 0.0159, p = 0.0529) **h**. Relative frequencies of task ordered and disordered concatenated sequences (ordered difference from 62.04%: chance level, Wilcoxon signed-rank; p =0.0215. Difference between groups, permutation test: observed difference in means = 50.3%, 99^th^ percentile permuted difference = 29.2%, p < 0.001). **i**. Example replay sequences with varied warp characteristics. **j**. Distribution of warp factors for forwards and backwards replay events (1x represents awake speed) (forward reverse difference, Wilcoxon rank-sum test, p = 0.4810). **k**. Normalised percentages of task and non-task related replay observed (task related difference from chance level: 50%, one sample t-test; p < 0.001. Difference between groups, Permutation test: observed difference in means = 21.20%, 99^th^ percentile permuted difference = 15.88%, p = 0.0004). **l**. Average (mean) replay event lengths observed for each recording session. **m**. Relative individual neuron involvement frequencies for each sequence during awake task activity and sleep periods (Exponential function fit by non-linear least squares, p < 0.001). **n**. Average (mean) start and end points for all forward (top) and reverse (bottom) replay events. Position is relative to the corresponding average awake sequence. **o**. Main: reactivation rates for each analysed sleep epoch against time from first sleep onset (OLS regression, r = -0.380, r^2^ = 0.145, p < 0.001). Inset: rate change against starting rate for each pair of analysed epochs per session. (for panels e-i, n = 19 sessions, n = 8 mice).

As with hippocampal-replay, during striatal-replay neural sequences often progressed faster than real world speed with the time course of the sequence being compressed (Fig.4i,j). Although neural sequences were on average more likely to occur faster than they were in the awake recordings, a lot of the sequences also occurred at real world speeds and could even progress slower. As with replay detected in other structures the individual reactivated neural sequences occurred in both forward and reverse direction (Fig. 4i,j). Across all events, roughly equal proportions of forward and reverse replay were observed, with replay speeds ranging from more than 5 times slower up to 20 times faster than awake activity. Task related neural sequences were more likely to be reactivated in post task sleep compared to non-task related sequences (Fig.4k) suggesting that the task related sequences were prioritised for reactivation. Also, neurons that were more frequently involved in replay events tended to be those which consistently participated in the same neural sequence during awake activity (Fig.4m). This relationship was non-linear such that the neurons that most consistently contributed to the awake neural sequence were more likely to occur in replay and the reverse was true for neurons that inconsistently contributed to the awake neural sequence. This suggests that on the single neuron level cells that consistently contributed to the awake sequence were prioritised for replay.

Like replay detected in the hippocampus our individual sequential events lasted around 100 – 400ms (Fig.4l)^36^. Analysis of the composition of each reactivated neural sequence revealed that replay tended to preferentially involve neurons which made up the central portion, rather than the boundaries, of each task related neural sequence (Fig.4n). The rate of replay also tended to decay from sleep onset and there was a significant relationship between time from first sleep onset and replay rate (Fig.4o). Analysing decay rate compared to current replay rate we found a strong relationship suggesting replay rate had nonlinear decay towards an equilibrium rate. Finally, analysis of recordings done during early learning revealed that replay characteristics were independent of learning stage (Extended Data Fig.9).

In summary, applying PP-Seq to post task sleep revealed that neural sequences that occurred during the awake experience were replayed during sleep. The features of this replay were consistent with replay in other areas such as the hippocampus, in that it occurred at time compressed and real-world speeds and occurred in forward and reverse order. The striatal replay appeared to be structured in a compositional manner such that individual neural sequences could be replayed individually, in awake order or even in combinations that rarely occurred during behaviour. The replay was also prioritised both at the level of the type of neural sequence and even at the level of individual neurons within a sequence, consistent with the idea that replay is important for the consolidation of both sequential order and refining the execution of individual parts of the behavioural sequence.

### Procedural memory formation and replay are independent of the hippocampus

Having established that task related neural activity is reactivated offline and shown that it shares many features with previously observed hippocampal replay, we next aimed to investigate whether these similarities were indicative of shared, mechanistic dependency. Indeed, even for ultimately hippocampus independent memories, initial consolidation is thought to be organised by hippocampal dynamics^1,2,16^. Hippocampal SWR events have been shown to display consolidation dependent coupling with replay in cortical areas^16,43^, suggesting hippocampal dynamics may be an essential trigger or driver of replay in other regions. To test whether the hippocampus is required for procedural memory formation in our task we performed large bilateral lesions to the hippocampus via injection of viral caspase across the extent of the hippocampus (see methods) (Fig.5a and Extended Data Fig.10a-d). Despite these large lesions, hippocampus ablated mice showed no learning deficits for the task when compared to controls. Lesioned mice reached the final task level in an equivalent number of trials and their learning curves were not significantly different from controls (Fig.5b). After learning (trials 4000 to 5000) mice completed the task with comparable movement speeds (Extended Data Fig.10e) and made a similar number of port-to-port transition errors (Extended Data Fig.10f). Also, like control animals, lesioned mice were highly task focused; rarely poking into task irrelevant ports (Extended Data Fig.10g). In sum, we find that large bilateral lesions to the hippocampus did not impair learning or expert execution of the sequence task. Lesioned mice were indistinguishable from controls across all measures of task performance.

**Figure 5:**
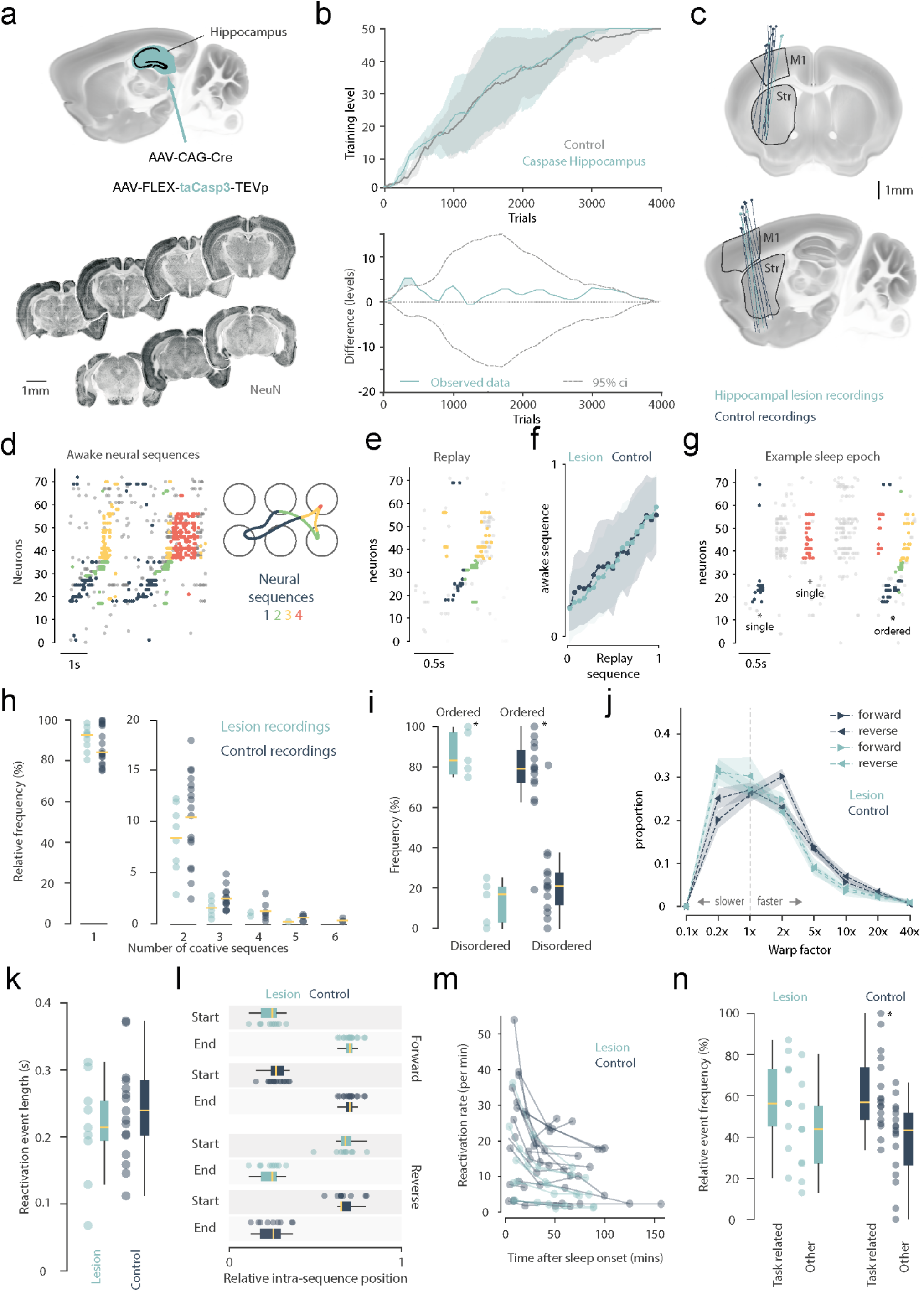
Procedural memory formation and replay are independent of the hippocampus. **a**. Top: schematic of the experimental approach for bilateral lesion of the hippocampus. Bottom: Histology, example NeuN stained coronal slices showing lesion extent across hippocampal volume for a mouse. **b**. Top: learning progression curves (training level vs trials) for control and lesion animal groups (shaded area denotes standard deviation). Bottom: differences in performance between the groups. Dotted lines indicate the 95% confidence interval for the shuffled data (see methods) (lesion n = 7 mice, control n = 6 mice). **c**. Schematic showing implanted neuropixel probe locations projected onto standard Allen atlas coronal (top) and sagittal (bottom). **d**. PP-Seq labelled sequences for example task related awake spikes. **e**. Example replay (same neurons as in d) of concatenated sequences that are ordered with respect to task related activity. **f**. Mean relative spiking positions in awake sequences vs. replay sequences for all analysed neurons across every session for lesion and control animals (Multivariate comparison between groups, MANOVA; Wilks lambda, p = 0.1797). **g**. Example sleep period with PP-Seq labelled replay sequences. **h**. Frequency of single (isolated) and concatenated events for each recording session (Differences between all groups, Kruskal-Wallis; p<0.001. Paired differences between lesion and control for each group, left to right, post-hoc Dunn test; p = 0.574, p =0.680, p=0.512, p=0.902, p=0.842). **i**. Relative frequencies of task ordered and disordered concatenated sequences with respect to observed awake task ordering for recordings containing 4 or more task related sequences (see methods). (Lesion, ordered difference from 62.04%: chance level, Wilcoxon signed-rank; p =0.0625. Control, ordered difference from 62.04%: chance level, Wilcoxon signed-rank; p =0.0215. Lesion, difference between subgroups, permutation test: observed difference in means = 73.9%, 99^th^ percentile permuted difference = 52.2%, p < 0.00386. Control, difference between subgroups, permutation test: observed difference in means = 50.3%, 99^th^ percentile permuted difference = 29.2%, p < 0.001. Pairwise difference for lesion and control ordered sequences, MannWhitney U test; p = 0.218). **j**. Distribution of warp factors for forwards and backwards replay events (1x represents awake speed. Differences for all warp factors, Kruskal-Wallis; p <0.0001. Paired difference for lesion and control, forward warp factor groups left to right, post-hoc Dunn test; p = 1.0, p = 0.0743, p = 0.93175, p = 0.712, p = 0.319, p = 0.489, p = 0.939, p = 0.748. Paired differences for lesion and control, reversed warp factor groups left to right, post-hoc Dunn test; p = 1.0, p = 0.210, p = 0.985, p = 0.971, p = 0.359, p = 0.562, p = 0.275, p = 0.987). **k**. Average (mean) replay event lengths observed for each recording session. (Independent t-test, p = 0.5089). **l**. Average (mean) start and end points for all forward (top) and reverse (bottom) replay events. Position is relative to the corresponding average awake sequence (comparison between all groups, ANOVA; p <0.001. Pairwise comparison between forward start groups, Tukey HSD; p = 0.749. Pairwise comparison between forward end groups, Tukey HSD; p = 0.949. Pairwise comparison between reverse start groups, Tukey HSD; p = 0.977. Pairwise comparison between reverse end groups, Tukey HSD; p = 0.880). **m**. Reactivation rates for each analysed sleep epoch against time from first sleep onset (Multivariate comparison between groups, MANOVA; Wilks lambda, p = 0.8961). **n**. Normalised percentages of task and non-task related replay observed (lesion, task related difference from chance level: 50%, one sample t-test; p = 0.289. Control, task related difference from chance level: 50%, one sample t-test; p < 0.0001. Lesion, difference between subgroups, Permutation test: observed difference in means = 16.9%, 99^th^ percentile permuted difference = 25.2%, p = 0.0626. Control, difference between subgroups, Permutation test: observed difference in means = 21.20%, 99^th^ percentile permuted difference = 15.88%, p = 0.0004. Pairwise difference for lesion and control task related sequences; independent t-test, p = 0.898). (Control: n = 19 sessions, n = 8 mice, lesion: n = 9 sessions, n = 3 mice).

Given procedural consolidation is independent of the hippocampus, a compelling hypothesis is that the hippocampus is not involved in shaping replay of procedural activity in the striatum. To test this, we performed large bilateral lesions of the hippocampus and then recorded neural activity in the DLS using implanted neuropixel probes (Fig.5c). Just like for baseline recordings (non-lesioned mice), applying PP-Seq to post-task sleep epochs revealed awake-like neural sequences replayed offline (Fig.5d,e). Across all measures, we did not find any differences in the characteristics of these events after hippocampus lesions. As observed in control recordings neural sequences were also ordered with respect to awake sequences at the single neuron level (Fig.5f). At a macro scale, sequences were also still replayed individually and in combination (Fig.5g,h). When neural sequences were replayed in combination, they were still more likely to occur in the order they occurred during the behavioural sequence (Fig.5i). The distribution of forward and reversed replay as well as the proportion of stretched or time compressed replay were the same between lesioned and control recordings (Fig.5j). There was also no difference in the rate, length, extent or decay of replay events between groups (Fig.5k, l, m and Extended Data Fig.11a). The proportion of task related replay was highly similar (Fig.5n) and analysis of single neuron contribution rates from awake activity to replay revealed no significant differences between groups (Extended Data Fig.11b). Together this suggests that the hippocampus has no role in generating striatal replay, in temporally compressing the replay, in compositionally ordering the replay, or in prioritising the replay at a neural sequence type or individual neuron level.

## Discussion

Our results demonstrate sequences related to procedural experience are replayed in the dorsal striatum during sleep. As with replay in other areas such as the hippocampus, the procedural replay we identify occurred at both real-world and time-compressed speeds^39^. Task related sequences were also preferentially replayed, and our unsupervised method revealed that this prioritisation also occurred at the level of individual neurons.

Surprisingly all these features were generated independently of hippocampus.

Previously it was proposed that the hippocampus may be critical for the active consolidation of both hippocampal-dependant declarative memory as well as memories that do not require the hippocampus for expression, such as procedural memory^2,3,11,17,44^. It was proposed that replay of a spatiotemporal context in the hippocampus could trigger the active consolidation of all kinds of memory by boosting context-related representations throughout cortical and subcortical circuits^3,11,17^. In contrast to this theory our results show that the hippocampus is not involved in either the generation of replay in the dorsal striatum or the consolidation of our procedural memory task. Our results support the alternative idea that there are “parallel memory systems”^23,24,45^ where consolidation can occur independently. We suggest that within the parallel memory systems replay may be a common mechanism for active consolidation. This suggests that either there are different sources of replay for distinct types of memory or that there may be a common unappreciated source for generating replay that is independent of the hippocampus. While our results support a body of literature that has shown procedural memories can be formed independently of the hippocampus^19,20,23-26,46,47^, others have shown that the hippocampus can in specific situations support the sleep-dependant consolidation of procedural memory^9,14,16,48-50^. In these latter cases it may be when the spatiotemporal context, or some other hippocampal dependant computation is important for learning^48^. Together our results support the theory that declarative and non-declarative memories can be consolidated in parallel but show that replay may be a common mechanism for active consolidation of both types of memory^23,47,51^.

If replay is a universal mechanism for memory consolidation, then what is the mechanism for driving replay if it is not the hippocampus? One option is that the hippocampus still generates replay for declarative memories and there is a different source for replay in circuits related to procedural and other types of non-declarative memory. However, even in cortical-hippocampal loops there is some evidence that patterned activity in the cortex precedes and predicts the content of replay in the hippocampus^27,28,52,53^. This suggests that replay may originate in parallel in both the cortex and the hippocampus. We suggest that it needs to be considered that replay is a phenomenon that occurs spontaneously in cortical and potentially other networks and that there might not be a single source.

### Unsupervised methods may help uncover the source of replay

Whether there are specific circuits that generate replay or not we propose that the use of unsupervised methods such as the one that we adapted here will aid in the investigation of replay. One advantage of these approaches is that they reduce the a priori assumptions about how replay should progress, such as that sequences of activity progress in a linear manner (for discussion see^35,39^). These assumptions have been shown to limit the types of replay that is identified. When recent methods that removed the assumption about the constant speed of replay were developed, hippocampal replay was found to progress at a range of speeds including real-world speeds^30^, just like the striatal replay we identify in this paper. Unsupervised approaches can also be used to examine replay without the generation of a behavioural template^54,55^, the weakness of this is that these approaches can identify neural sequences that are not related to the behavioural features of interest. This might make these approaches better suited for tasks with repetitive task structure, like our task. When these methods are applicable we show that they can in practice be more sensitive and accurate than even state of the art decoding methods.

### Striatal replay

Our discovery that replay occurs in the dorsal striatum independently of the hippocampus is consistent with previous work that has shown that there is reactivation of task related ensembles in the dorsal striatum following motor skill learning and that this occurs in combination with an increased synchrony between cortical and striatal ensembles^33,56^. We now show that the features of this reactivation mirror those associated with replay in the hippocampus and cortex. This sets the stage for investigating the role replay plays in a host of dorsal striatum dependent tasks from, perceptual learning to value based decision-making^57-60^. It will be of particular interest to determine how striatal replay is coordinated with dopamine release as both phenomena are strongly linked to memory formation^57,59,60^. Indeed, in quiet wakefulness reward related neurons in the ventral tegmental area, putatively dopamine neurons, are coordinated with hippocampal replay but this is not the case during sleep^61^.

Despite this, dopamine has been shown to have a causal role in offline memory consolidation^62-65^ and whether this is due to coordination between replay and dopamine release in the dorsal striatum needs to be investigated.

### Limitations

Currently we have shown that offline processing in the striatum is needed for procedural memory consolidation and suggest that replay might be the process that drives this active consolidation. To prove this, it will be important to selectively disrupt striatal replay and assess the impact on memory formation. A similar approach is possible in the hippocampus because replay is believed to occur almost exclusively during sharp wave ripples^40^. Due to this, closed-loop inactivation of the hippocampus during ripples has been used as a good proxy for disrupting replay and showing it causally contributes to consolidation^66-69^. Without a biomarker for striatal replay a similar approach will not be possible, new approaches may need to be developed to rapidly detect the onset of replay to target these patterns for disruption.

### Summary

In conclusion we have shown that replay occurs in the dorsal striatum following procedural learning and have demonstrated that this process is independent of the hippocampus. We propose that replay is a common mechanism used to actively consolidate all forms of memory during sleep, but that source of replay may be distinct depending on the type of memory. Future work will be needed to identify the sources of hippocampal independent replay and determine how replay in distinct networks is coordinated for the appropriate consolidation of memory.

## Materials and methods

### Animals

Adult male and female Mice (8-50 weeks) C57BL/6J (wild-type) were used. Mice were housed in HVC Cages with free access to chow and water on a 12:12 h inverted light:dark cycle and tested during the dark phase. Mice used in sleep recording experiments were housed on a non-inverted light cycle and tested during the light phase (normal daylight hours). For behavioural experiments, mice were water deprived. Animals had access to water during each training session, and otherwise water was administered by hand. Water was supplemented as needed if the weight of the mouse was below 85%. All experiments were performed in accordance with the UK Home Office regulations Animal (Scientific Procedures) Act 1986 and the Animal Welfare and Ethical Review Body (AWERB). Animals in test and control groups were randomly selected.

### Behavioural procedures

Mice were trained in custom built behavioural arenas measuring approximately 16cm x 19cm x 24cm (width, length, height). Box walls were made of 0.5cm thick opaque white or transparent red acrylic and had 8 poke ports mounted on the front wall. Ports (sanworks, ID 1010) protruded 2cm from the wall into the area and were arranged in a 4 x 2 grid such that neighbouring ports (vertical and horizontal) were 3cm apart (centroid to centroid). Ports contained side mounted infrared LED and sensor to detect poke events and a back mounted visible light LED to illuminate the port. Each port also contained a waterspout for reward delivery. Poke events were registered by Sanworks port PCBs (ID: 1004) connected to a Bpod (ID: 1027) programmed with a custom behavioural protocol (MATLAB). 2ul Water delivery was triggered by Bpod via a connected Miniature Solenoid (Lee Company: LHDB0513418H). Sounds were played at port entry via a speaker (DigiKey part number: HPD-40N16PET00-32-ND) and amplifiers (DigiKey part number: 668-1621-ND).

Mice were rewarded for completing the full 5 step poke sequence. No reward was given if animals missed a step in the sequence, but animals were not punished for adding extra steps into the sequence. Single trial events were defined as all poking activity that led to reward delivery at the final port. Within a single trial, animals could make multiple attempts at completing the sequence or add additional elements to the sequence and still eventually receive reward. The task was self-paced though if an animal initiated a trial but did not register a poke into any port for 30s this trial timed out, was left unrewarded, and a new trial was cued. Across all levels, when animals entered a port (breaking the IR beam) a short duration sibilant noise was played.

Poke sequences were shaped by an automatic protocol with 50 training levels of predefined difficulty. Mice started from the lowest level (Level 1) and progressed up to the final task (Level 50) (Extended Data Fig.1a-b). During training, performance was assessed every 10 trials and this metric determined whether mice progressed up a level (performance > 90%), regressed down a level (performance < 20%), or remained at the same training level (20% < performance < 90%). In Early task levels (1 – 12) mice were rewarded for performing each step of the 5-step sequence. With progression to higher levels, reward steadily decreased and then switched off port-by-port until reward was only given upon reaching the final port (levels 12 - 50). In early task levels (1-12) each step in the sequence was also visually guided by successively illuminating port LEDs which were switched off port-by-port once the animal had successfully poked. After this (levels 12 – 49), across successive levels LEDs brightness was gradually dimmed port-by-port and eventually turned off permanently, except for the initial port in the sequence which remained illuminated at the start of each trial to signal a new trial was available. At the final stage of the task (level 50) only the first port in the sequence was illuminated in this way. Even after reaching the final task (level 50), during training animals could drop down to lower levels if they performed badly. However, in circumstances where it was necessary to test animal performance at the full task, mice were held at level 50 for the duration of the session. For AP5 infusion experiments, to increase sensitivity to performance changes during test sessions, training performance was assessed (and training level updated) every 4 trials. For hippocampus lesion experiments animals were trained on the task for at least 5000 trials except for one animal which was only trained for 2004 trials. This animal was excluded from analysis of post learning performance (Extended Data Fig.e-g) for this reason.

### Surgical procedures

Mice were anaesthetised with Isoflurane (0.5–2.5% in oxygen, 1 l/min) - also used to maintain anaesthesia. Carpofen (5 mg/kg) was administered subcutaneously before the procedure. Craniotomies were made using a 1-mm dental drill (Meisinger, HP 310 104 001 001 004). Injections were delivered using pulled glass pipettes (Drummond, 3.5”) on a stereotaxic frame (Leica, Angle TwoTM).

For striatal lesions, initial lesions were excitotoxic via injection of NMDA (2mg/100mL), though for most animals shown lesions were achieved by injecting a 1:1 mix of AAV2/1-hSyn-Cre (1014 vg/ml) and AAV2/5-EF1a-DIO-taCasp3-T2A-TEVp (1014 vg/ml) as this proved more successful. The mix was diluted 5 times in saline buffer prior to injection. For control animals, saline or GFP virus AAV2/5-CAG-EGFP (1043 vg/ml) was injected instead. In each hemisphere 4 injections (∼80nl each) at 3 different depths were made to distribute the viruses as evenly as possible and to provide enough coverage, injections were targeted to the DLS or DMS dependent on the experimental group. For DLS, injections were made at coordinates AP: 0.2 to 0.8mm ML: 2.5 to 2.7mm DV: -3.0mm to -3.7mm (where a range is given, injections were given at regular spacing between these values). For DMS, insertions were made at coordinates AP: 0.2 to 0.8mm ML: 1.8mm DV: -3.0mm to - 3.7mm.

For hippocampus lesions, this same cre-caspase mixture as used in the striatal lesion experiments was injected. Control animals underwent the same surgical procedures except saline was injected instead. Injections were made bilaterally at 13 locations per hemisphere (see Table 1). After surgery, animals were given at least 3 weeks of recovery before training was started.

**Table 1:**
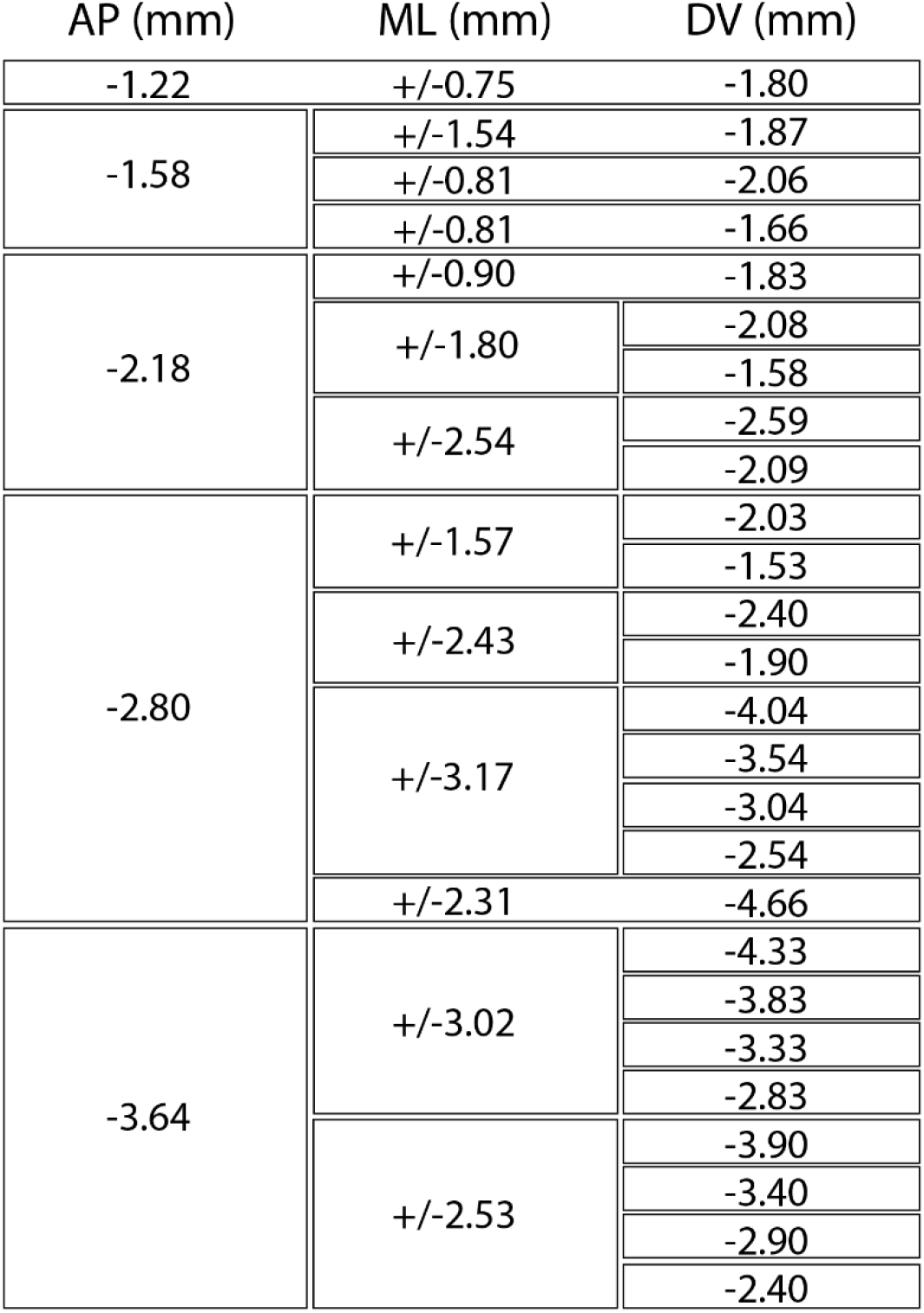
Positions of injections into hippocampus.

For Cannulation experiments, 5mm 26-Gauge cannulas (P1 technologies, cat number: C315GS-5/SPC) were implanted at coordinates anterior posterior (AP) 0.5mm, medial lateral (ML) 2.9mm and dorsal ventral (DV) 2.0mm from pial surface, with a 10 degrees’ tilt (tip tilted towards midline). For optogenetic experiments, we implanted flat optical fibers of 200μm diameter (Newdoon: FOC-C-200-1.25-0.37-7) at AP - 1.6mm, ML -1.9mm, DV 2.8mm (20 degrees’ tilt). Implants were affixed using light-cured dental cement (3m Espe Relyx U200) and dental cement (Super-Bond C&B Bulk-mix, Sun Medical) and the wound sutured (6-0, Vicryl Rapide).

For neuropixel implantation, prior to training animals underwent an initial surgery where the skull was exposed, coated with a thin layer of dental cement, and marked for later skull levelling. A craniotomy was drilled over an arbitrarily chosen region of posterior cortex and a ground pin was implanted. To replicate the weight and size of the eventual implant, mice were trained on the task with a size and weight matched dummy implant, this was fixed during initial surgery to the cement layer using silicon (Kwik-sil: World Precision Instruments). For probe implantation, the dummy implant was removed, and the skull was levelled using previously noted skull markings. A craniotomy and durotomy was made at coordinates AP 0.8mm, ML = 2.1mm and the probe was implanted to a depth of 4.0mm at a 10-degree angle (tip tilted away from midline). The external grounding wire was fixed to the previously implanted skull pin. The craniotomy was then sealed using Duragel (Cambridge Neurotech) and a 3D printed casing was fitted around the probe for protection. Implants were performed using a retrievable system and, in some cases, animals were reimplanted with a second probe. At the end of the experiment, probes were recovered for future reuse.

### Electrophysiological recordings

Animals were habituated to the size and weight of the implant by first training on the task with a size and weight matched dummy implant fixed to the skull. Dummy implants were constructed from a 3D printed plastic casing, aluminium implant cassette and surgical tape. During training, to simulate eventual recording conditions animals were tethered to an overhead cable connected to a motorised rotary joint (Doric, B330-1027-001). Mice were also habituated to sleeping in their home cage while connected to this tether. To increase sleep incidence during recordings, all mice (except for one) were housed with a normal light dark cycle. Neuropixel 1.0 (phase3B) probes were implanted through motor cortex and striatum and after implantation, daily recording sessions were conducted as continuous recordings across sleep and task epochs (3-6 hours).

Recordings were acquired using neuropixels acquisition hardware (imec neuropixels 1.0 headstage, interface cable and PXIe Acquisition Module) with open-Ephys software. Post-acquisition spike sorting was done using Kilosort3^70^. Spike sorting was checked using Phy2 (https://github.com/cortex-lab/phy/) but analysis was not performed on manually curated clusters.

### Pharmacological manipulations

For Muscimol infusions, ∼30nL of either muscimol (Sigma-Aldrich) at 0.2mg/ml or saline buffer were infused via implanted cannulas. The infusion system consisted of a 1µl Hamilton syringe (Merck), plastic tubing (P1 technologies cat no. 8F023X050P01) and injection cannulas (P1 technologies cat no. C200IS-5/SPC). Tubing was filled with mineral oil to ensure an air-tight setup for accurate volume administration. Animals were briefly headfixed and infused at a rate of 10nl/min for 5 minutes per cannula. Animals were then allowed to rest in their home cage for 10-15 minutes and tested on the task. Between muscimol infusion experiments animals were given recovery break of at least a day and task behaviour was assessed on this break day ensure performance returned to that pre-infusion. All animals were habituated to headfixing prior to experiment onset.

For AP5 experiments, we adapted methods described previously^33^. We bilaterally infused 450nl saline or 450nl of 5µg/µl D-AP5 (Bio-Techne, diluted in saline), via cannulae implanted into the dorsolateral striatum. Immediately after training, animals were headfixed and infusions were carried out at a rate of 65-90 nl/min for 5-7 minutes per cannula. After infusion animals were returned to their home cage. In the test session the next day (approximately 24 hours later), animals were trained on a performance sensitive version of the behavioural task (see Behavioural training). This allowed for higher sensitivity in detecting changes to task performance. Infusions during learning were done from levels 12 to 49 (after the reward guided habituation phase: levels 1 to 11). Infusions were done if animals climbed at least 3 levels that session but were not done on consecutive days. Before the experiment, animals were habituated to head-fixing. Infusions of saline and AP5 were alternated throughout the learning curve of each animal. After learning, once animals reached stable expert performance (level 50) for at least 4 days, infusions of either saline or AP5 were given for 4 consecutive days. All mice were used for both experimental groups and so before switching the type of infusion animals were trained until at least 4 days of stable expert performance was seen. One animal was unable to reach level 50 with stable performance so was excluded from this experiment.

### Tissue processing and image analysis

At the end of experiments, animals were euthanized via intraperitoneal (IP) injection (10 ml/kg pentobarbital) and brain tissue fixed via vascular perfusion (4% paraformaldehyde) and collected for histology. Prior to implantation, neuropixel probes were coated in dye (DiI) for later visualisation. Brains were imaged using a serial section two-photon^71^.Our microscope was controlled by ScanImage Basic (Vidrio Technologies, USA) using BakingTray, a custom software wrapper for setting up the imaging parameters:

§ https://github.com/SainsburyWellcomeCentre/BakingTray,https://doi.org/10.5281/zenodo.3631609

Images were assembled using StitchIt:

§ https://github.com/SainsburyWellcomeCentre/StitchIt,https://zenodo.org/badge/latestdoi/57851444

The 3D coordinates of the injections, fiber, cannula and neuropixel probe placements were determined by aligning brains to the Allen Reference Atlas (Allen Reference Atlas – Mouse Brain. Available from atlas.brain-map.org.) using brainreg^72^, and the tracks were located and traced in common atlas coordinates (brain-reg segment).

Brain slices were all stained following the same procedure: Blocking in staining solution (PBS + 1%BSA + 0.5%Triton-X) for 1 hour. Primary antibody(s) (1:1000 in staining solution) for 2-4 hours at room temperature or overnight at 4 degrees Celsius while rocking. 15 minutes wash with staining solution. Second antibody(s) (1:1000 in staining solution) and DAPI for 2 hours at room temperature while rocking. Slices were then washed in PBS and mounted using Prolong or ProGold mounting medium (Thermo-Fischer). Primary antibodies used were NeuN (abcam, ab177487) & GFAP (abcam). Secondary antibodies used were Alexa-488 anti-chicken (Thermo Fischer, A-11039) and Alexa-647 anti-rabbit (Thermo Fischer, A-21245).

For striatal lesions brains were sliced using a cryotome at a thickness of 40um and with a vibratome at 100um for hippocampal lesions. 15 to 20 slices covering the entire region at regular intervals were selected for NeuN and GFAP staining. Slices were mounted in standard glass slides using standard mounting medium, and subsequently imaged in the Slide Scanner (Zeiss) using a 20x objective. Lateral and medial striatum were defined by the extent of axons from prelimbic/cingulate projections and motor cortical projections respectively (Allen projection experiment numbers: 157711748, 112514202, 180720175 & 180709942). The lesioned areas for each of these regions was determined manually for each slice using Brainreg segment. For striatal lesions mice with lesions that had more than 20% of volume overlapping with cortex were excluded from analysis.

### Performance measures

A trial was defined as all the poke events that occurred proceeding reward or trial time out (no pokes for 30s) Performance was calculated per trial and involved segmenting sequence pokes into ‘attempts’ which were temporally relevant (within 2s of each other). Attempts were considered as starting only from the initiation port (port 1), any pokes that occurred before the first poke into port 1 were excluded. An attempt was assigned a value of 1 if it contained the perfect poke sequence and 0 if it contained an error. The mean score across these attempts was then calculated. During training a simplified measure was used to calculate ongoing trial by trial task performance used for updating training level; rather than across ‘attempts’, the same measure was calculated across trials. Ongoing performance was scored as a mean over a window of 10 trials except for AP5 test sessions where this window was reduced to 4 trials.

### Video tracking analysis

Videos were captured at 60 fps and mouse movements were tracked using DeepLabCut^73^. Tracking points below 98% confidence interval were excluded and replaced by interpolating between accepted points. Task movement variability was first calculated individually for different task subsequence (movement vectors). To achieve this, only trajectories that passed close to each port in the subsequence (within a 1cm radius) in order and with appropriate timing (within a 2s port-to-port time window) were considered. Trajectories were averaged to find the mean trajectory curve. This curve was then segmented into 3000 spatial bins to be used as a reference and the distances between each data point in each trajectory and their closest spatial bin were noted. These distances were then used to calculate the standard deviation (movement variability) of each tracked trajectory from the mean curve. To create the standard space sequence occurrence plots during analysis of PP-Seq output data similar analysis was done. However, average trajectory curves were generated for the entire task sequence rather than individual subsequence chunks.

### PP-Seq replay detection

We adapted PP-Seq^41^ such that after training the model on a set of awake data, the parameters which defined each neural sequence could be fixed per their fit from the awake data. This allowed us to then apply each trained model to sleep data, in order to search for the same recapitulated neural sequences. Our adapted model can be found here:

https://github.com/lindermanlab/PPSeq.jl/pull/15.

Running PP-Seq^41^ required setting 12 hyperparameters. The three values related to background firing were set accordance with reported literature, as were those related to width of a neurons response. Of the seven remaining hyperparameters 5 were chosen by grid search (see Table 2) (Extended Data Fig.6), the other two were found in preliminary work to have little effect on the overall PP-Seq output. We used cross-validation to train and test the model: a subset of spikes was held-out from the data, the rest used to train the model, then the log-likelihood of the held-out spikes was measured under the trained model. The hyperparameters that led to the highest held out log-likelihood were taken as those which best capture the structure in the data and could predict held-out spikes. We chose our selected model from the top 20 models (all within error of each other) by visual inspection of the output labelling. Together this specified the 12 hyperparameter values that we used for our subsequent analysis. We performed this hyperparameter fitting on the data from one animal, then used the same values, occasionally scaled for the number of neurons and average firing rate in the data, for all other mice.

**Table 2:**
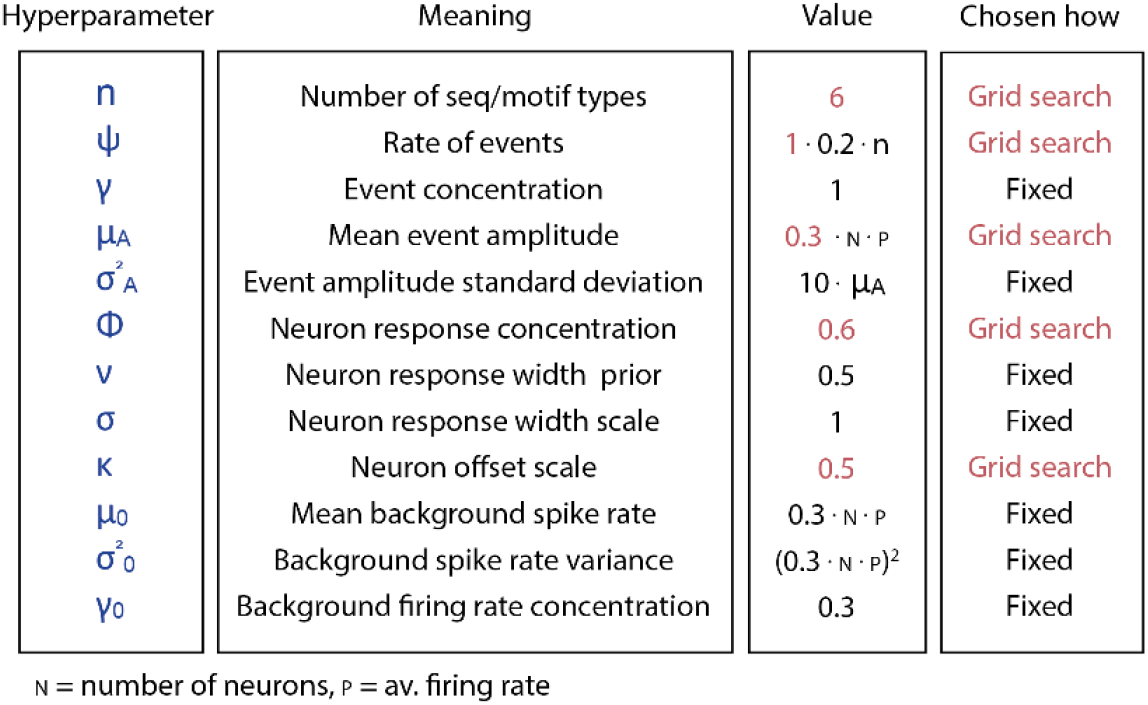
PPseq model hyperparameters.

The PP-Seq algorithm was then applied to each recording session individually. Striatal neurons were first filtered to remove high and low firing rate units (Fano factor 0.5 -12) and a 600s period of high task engagement awake activity was chosen to fit the model on. To sample replay during our offline recordings PP-Seq was applied to segments (500-1500s) of the recording totalling at least 2000s throughout the sleep period. When applying PP-seq during sleep we permitted each model to fit an additional 2 latent sequences and allowed a range of time warp values. Hence, after applying to sleep, up to latent 8 sequences, with various timewarps could be reported by the model.

After running PP-Seq, the neural sequences in awake and the sleep recordings were pre-processed before further analysis was performed. Since PP-Seq is a probailistic model its outputs represent a posterior over the assignments of the spikes to each latent, in the form of many samples from the posterior. We made use of this to filter for only high confidence assignments, by analysing only spikes that were consistently assigned to a sequence with 75% probability. Single replay events were defined by binning PP-Seq labelled spikes into 20ms time bins and grouping bins together if they were adjacent and contained spikes for a given sequence type. Replay instances were only analysed during classified sleep periods and replay events were then excluded from analysis if they did not contain at least 5 spikes. Coactive replay events were defined as any sequence of events that occurred within 300ms of each other. Replay events were filtered further by regression analysis of their progression with respect to average awake sequential position. Regression slopes were used to define warp factor for each replay event. For analysis of coactive sequence ordering, ordered coactive pairs of sequences were those which respected observed task ordering (forward or reversed) or pairs which were a repeat of the same sequence type. Disordered sequence pairs were those which did not normally occur adjacent to each other in the task sequence (forwards or reversed). Recordings with only 3 task related sequences were excluded from this analysis as they could not contain disordered pairs. To determine replay propagation extent compared to awake sequences, neuron-to-neuron spike ordering for each event was compared to the expected awake ordering for that event type.

### Bayesian decoders

The state space decoder (decoder 1) used was as described in^39^. Models were trained for each recording session to predict a two-dimensional posterior (video tracking position) from spiking data. The linear decoder (decoder 2) used was as described in^42^. Models were trained for each recording to predict a one-dimensional posterior (linearised task space taken from average video tracking position). Training data for each model was filtered to only include successful movement trajectories. Filtering was done spatially: only trajectories that passed close to each port (within a 1cm radius) in order and with appropriate timing (within a 2s port-to-port time window) were considered. Spatial bins were 3 times the average distance travelled in one-time bin (20ms), on average this corresponded to approximately 400 spatial bins.

Replay detection was done by applying the trained models to short data segments of interest; usually 1-5s of data. For decoder 2, detected events were defined as true replay or noise by quantifying the spatial coherence of the decoded position. This was defined by whether the number of spatial bins necessary to explain the prior position up to a 95% confidence interval was within a threshold value. Thresholds were calculated prior to applying the decoder to sleep data. For each event type in each recorded session the threshold number of bins was calculated from the distribution of 95% posterior density for more than 200 hundred awake events and periods of random noise activity. Based on these distributions, the threshold value was set to maximize the number of true positive while minimizing the number of false positive events.

For decoder 2, replay was assessed as described in^42^. Putative replay events were first scored using a line-fitting algorithm and events with a reasonable linear fit were then assessed for significance through a shuffle analysis. Both of these analyses depend on a width parameter which scales the degree of variation from linear fit which is accepted. This parameter was set dynamically for each model by comparing the rate of true and false positive synthetic replays found. The chosen parameter for each model was that which maximized true replay detection while minimising false positives.

### Synthetic data testing

The synthetic replay data was generated by implanting PP-Seq detected sequences into background noise. Background noise was generated by randomly permuting neuron IDs from awake activity. Hence, spiking content in background noise and the original spike data was identical, but the neuron Ids given to PP-Seq were shuffled. For each test, sequences were extracted from the corresponding PP-Seq labelled awake dataset by manually setting an inclusion zone generated by a time window centred on the middle of the detected sequences. All spikes in each inclusion zone were extracted. Sequences were then filtered for representative, non-contaminated sequences. This was done by excluding the top and bottom 25% of sequences based on total sequence spikes and the number of contaminant (other sequence) spikes in these windows. Sequences that did not occur regularly in the labelled data were excluded from this analysis. From all extracted sequences, 200 sequences were then chosen (selected to maximise equal numbers from all sequence types extracted) for implantation into 600s of noise. When required, sequences were then manipulated and altered. For warp values which stretched the sequences, fewer than 200 were implanted to avoid excessive overlap between sequences. In rare cases where less than 200 sequences were originally extracted in total then random sequences from underrepresented groups were duplicated to make up this number. The chosen sequences were then randomly ordered and implanted into noise roughly equally spaced apart. Noise data was deleted where sequences were implanted such that the implanted sequences replaced spikes in these regions. For PP-seq testing, replays were considered correctly labelled if they were identified as the same sequence type that was inserted in the synthetic data, otherwise they were deemed mislabelled. For decoder testing, events found that had a spatial overlap of at least 40% with the original inserted replay location were considered as correctly labelled events, those that didn’t were deemed mislabelled.

### Sleep state classification

For each analysed epoch, sleep state was determined using LFP and movement. LFP from several (minimum 3) evenly spaced striatal electrodes along the neuropixel was pre-processed by manually excluding noisy electrodes, z scoring, and then a mean striatal LFP signal was generated. Movement was determined from video tracking. Processed, z-scored tracking points were averaged and used to determine mean movement velocity for each time window. Movement velocity below a threshold (0.8 times Standard deviation) was classified as putative sleep. Putative sleep periods were then validated using delta and theta spectral power as has been described previously ^35^.

### Statistical analysis

A one sample t-test (1 group), paired t-test (2 groups) or one-sample ANOVA (3 or more groups) were carried out when the assumptions for a normal distribution of observation within groups (Shapiro-Wilks test) were satisfied. Otherwise, the non-parametric Wilcoxon signed-rank, Mann-Whitney U test or Kruskal-Wallace test were used. When there were unequal observations between groups, an independent t-test was used, given the assumptions were satisfied. When data (2 or more groups) were multivariate a MANOVA was performed. For the statistical analysis of training learning curves, the data was down sampled into bins of 100 trials. Each animal’s learning curve was randomly reassigned to the lesion or control group to look at the mean difference between controls and lesions. This shuffling of learning curves was repeated 10000 times, and all 10000 means were used to find the 95% confidence interval for the difference in the data due to chance. For comparison between task and non-task related replay sequences a similar permutation test was performed and repeated 10000 times. The resultant distribution of means was analysed and the 95% confidence interval reported. For analysis of coactive replay rates, expected percentages were calculated as the independent Poisson probability given the mean rate of replay events (individual sequences) and a time interval of 300ms plus average replay event length. For ordered and disordered replay analysis, expected percentage ordered and disordered were calculated from the expected ratios of ordered and disordered given the number of task related sequences in each recording. These expected ratios were summed across recordings to find overall expected percentages.

## Supplementary videos

[Supplemental videos link](https://liveuclac-my.sharepoint.com/:f:/g/personal/ucqfmrj_ucl_ac_uk/Enn3UpunLz9OvMOxoD1woSQBVf7Cxze3z4pOCA0X2LDZDA?e=F48C3t)

## Extended Data Figures

**Extended Data Figure 1:**
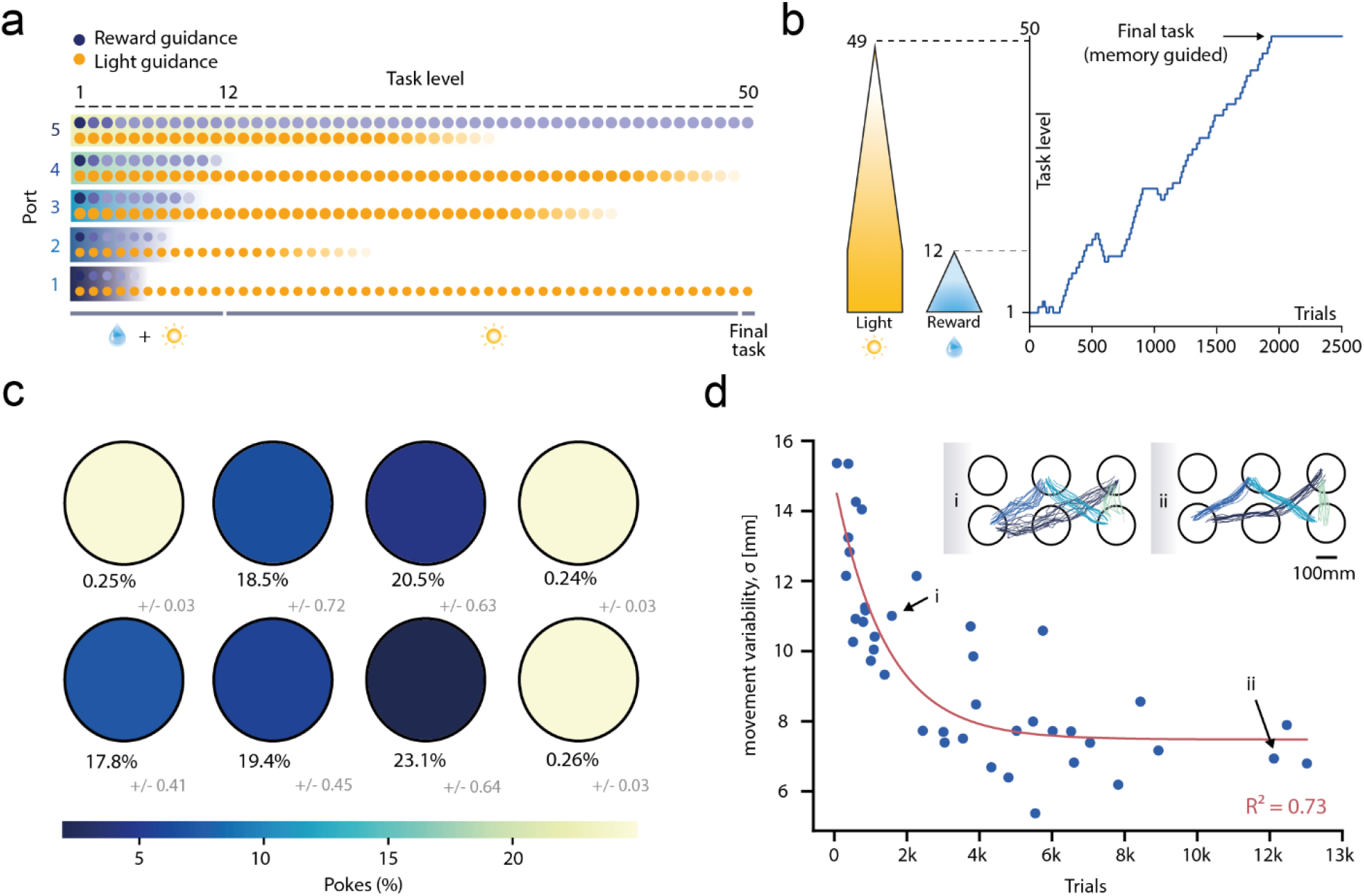
A novel sequence learning task based on sequence learning. **a**. Schematic showing light guidance and reward delivery at each port in the task sequence for each training level. **b**. Example animal training level progression and regression learning curve. Reward and light guidance shut off points are denoted by dotted lines. **c**. Port poke occurrences (mean) for expert animals (trials 3000 to 3500, n = 33, SEM for each port shown in grey). **d**. Average movement variability (standard deviation from average trajectory) across all subsequence task movements for mice at different levels of task experience (n = 8 mice). Above: example tracking data showing subsequence trajectories from a novice mouse (i) and an expert animal (ii). Number of trials is total completed prior the tracking session (tracking point was centre of the head).

**Extended Data Figure 2:**
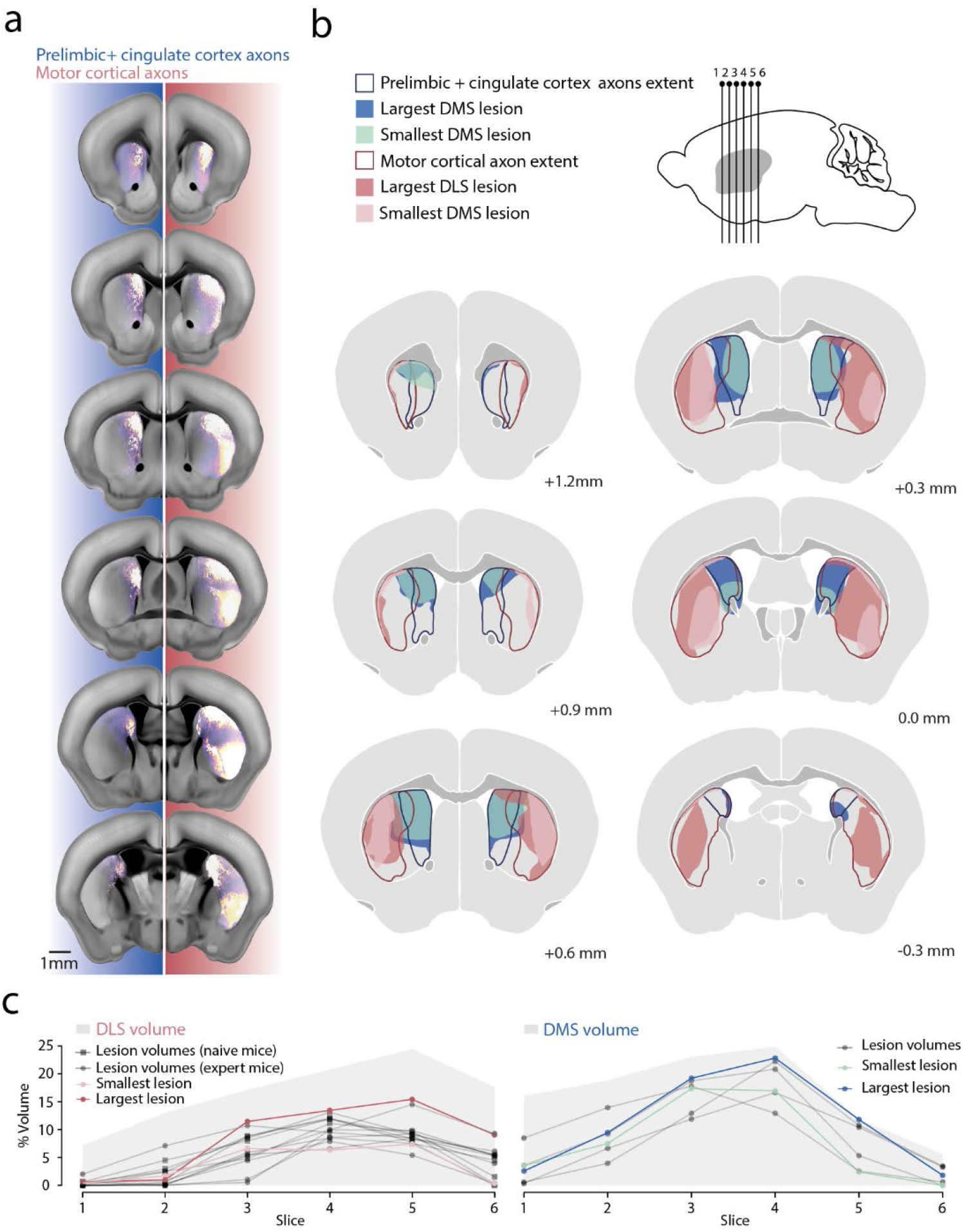
Striatum lesion extent. **a**. Allen reference projections from prelimbic and cingulate cortex (left, Allen experiment: 157711748 & 112514202) and motor cortex (right, Allen experiment: 180720175 & 180709942). **b**. Allen projection defined DMS (blue line) with largest DMS lesion (blue shaded) and smallest DMS lesion (turquoise shaded) for the extent of striatum. Allen projection defined DLS (red line) with largest DLS lesion (red shaded) and smallest DLS lesion (pink shaded) for the extent of striatum. **c**. Percentage of total DLS and DMS volumes lesioned for each animal in the two respective groups. Total volume of each region is shown (grey background) as well as the smallest and largest lesions in each group (light and dark coloured lines). DLS lesions are further subdivided into those animals lesioned while naive; for pre-training experiments (square markers), and animals that were lesioned while task experts for post-training experiments (circle markers).

**Extended Data Figure 3:**
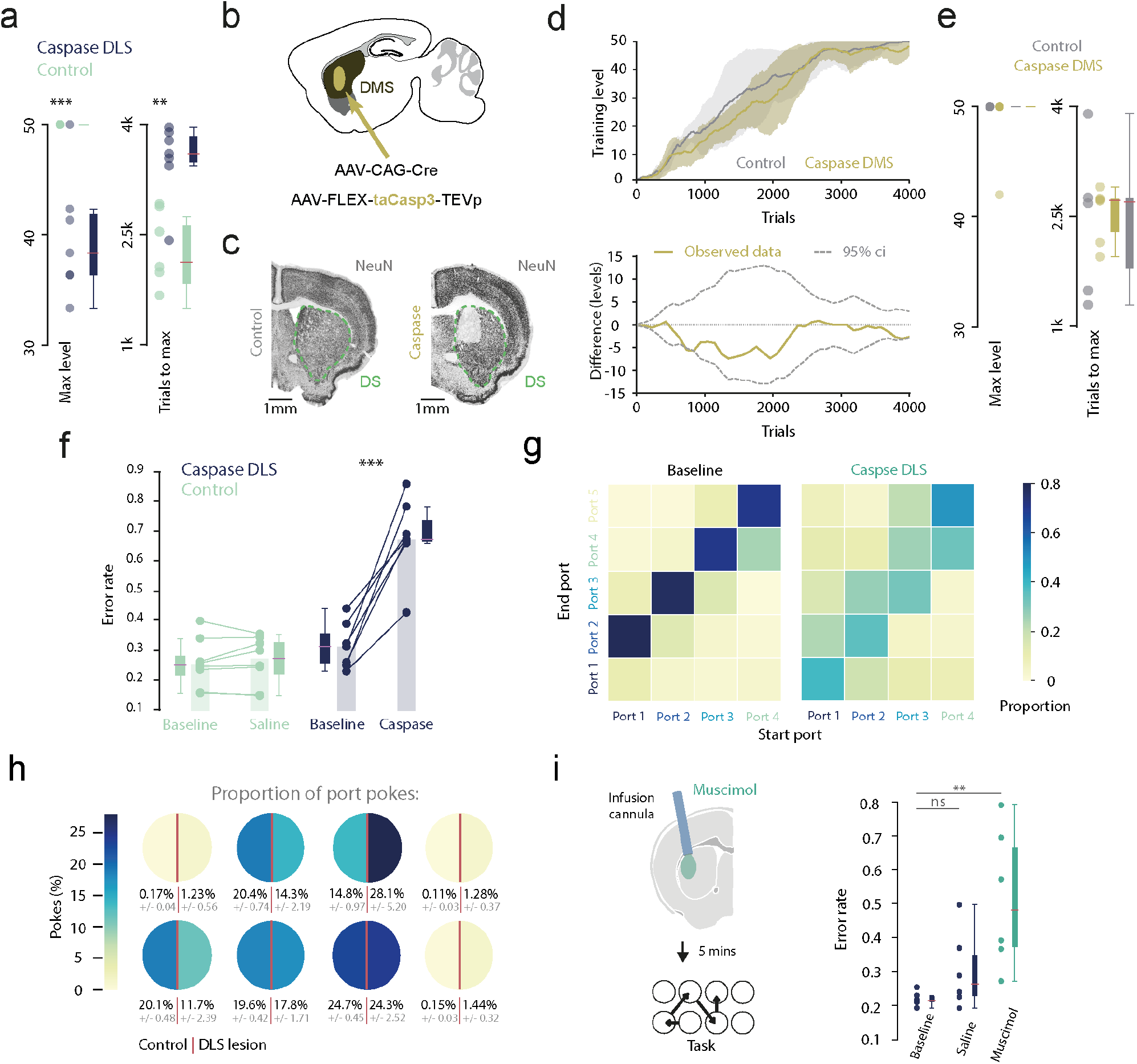
Task learning and execution are dependent on the DLS and not DMS. **a**. Left: maximum training level achieved within criterion (4000 trials) for DLS lesion and control animals. Right: trials to reach maximum attained level (max level, p = 0.001, trials to max, p = 0.0008, independent t-test, lesion; n = 7 mice, control; n = 6 mice). **b**. Schematic of the experimental approach for bilateral lesion of the DMS. **c**. Histology; coronal sections showing neurons in one example hemisphere of the striatum (green outline) for DMS lesion and control mice. **d**. Top: Learning rate (training level vs trials) progression curves for control and lesion animal groups (shaded area denotes standard deviation). Bottom: differences in performance between the groups. Dotted lines indicate the 95% confidence interval for the shuffled data (see methods). **e**. Left: maximum training level achieved within criterion (4000 trials) for DMS lesion and control animals. Right: trials to reach maximum attained level (DMS lesion; n = 6 mice, control; n = 6 mice). **f**. For DLS lesions and controls, proportion of port-to-port transition errors in the 3 sessions before and 3 sessions after injection surgery (p = 0.004, paired t-test). **g**. Average (mean) transition histograms before and after injection surgery. **h**. Average (mean) percentage port poke occurrences for all animals in control (left semi-circles) and DLS lesion (right semi-circles) groups for the 3 sessions after injection surgery (grey numbers are SEM). **i**. Left: schematic showing experimental design for muscimol infusions (bilateral). Right: port-to-port transition error rates for baseline sessions, saline infusions and muscimol infusions (ANOVA; p = 1.0e-5, Tukey HSD; p = 0.0018).

**Extended Data Figure 4:**
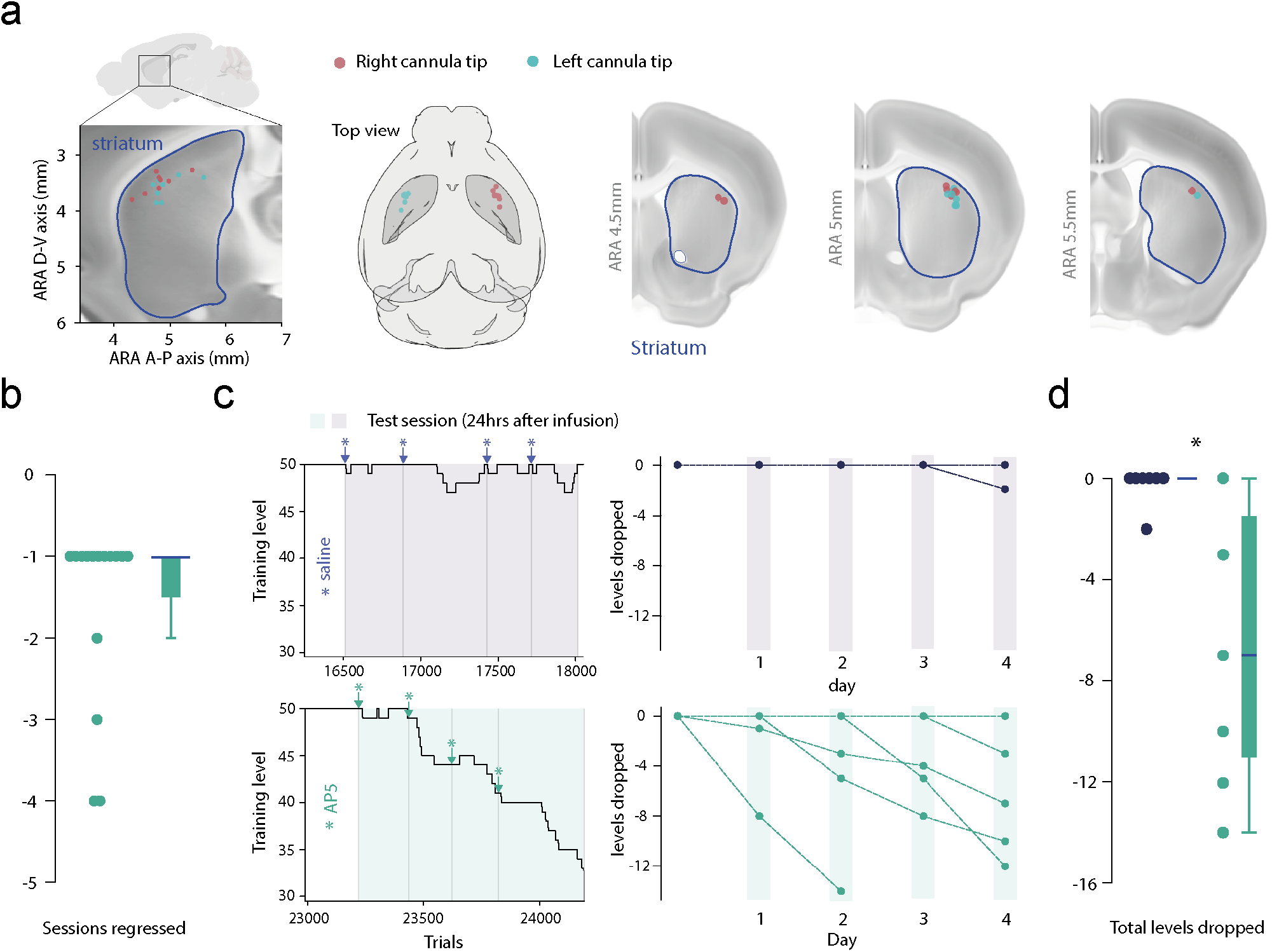
Blocking offline plasticity in the DLS impairs late stabilisation of procedural memory. **a** Left and right hemisphere cannula tip positions in Allen reference atlas coordinates. left to right: Sagittal view, top-down view, and coronal sections. **b** Number of sessions since minimum level was last seen for AP5 infusions experiments that resulted in negative trial changes. **c**. Left: example training level progression for fully trained animals given consecutive infusions (infusions points marked by stars, sessions are marked by grey vertical lines, shaded regions are test sessions – 24 hours post infusion). Right: cumulative levels dropped across the four consecutive infusion days for each animal. **d**. Summary plot showing total levels decreased for each infusion type (AP5 & saline) after 4 days of consecutive post session infusion (p = 0.024, MannWhitney U, n = 6 mice).

**Extended Data Figure 5:**
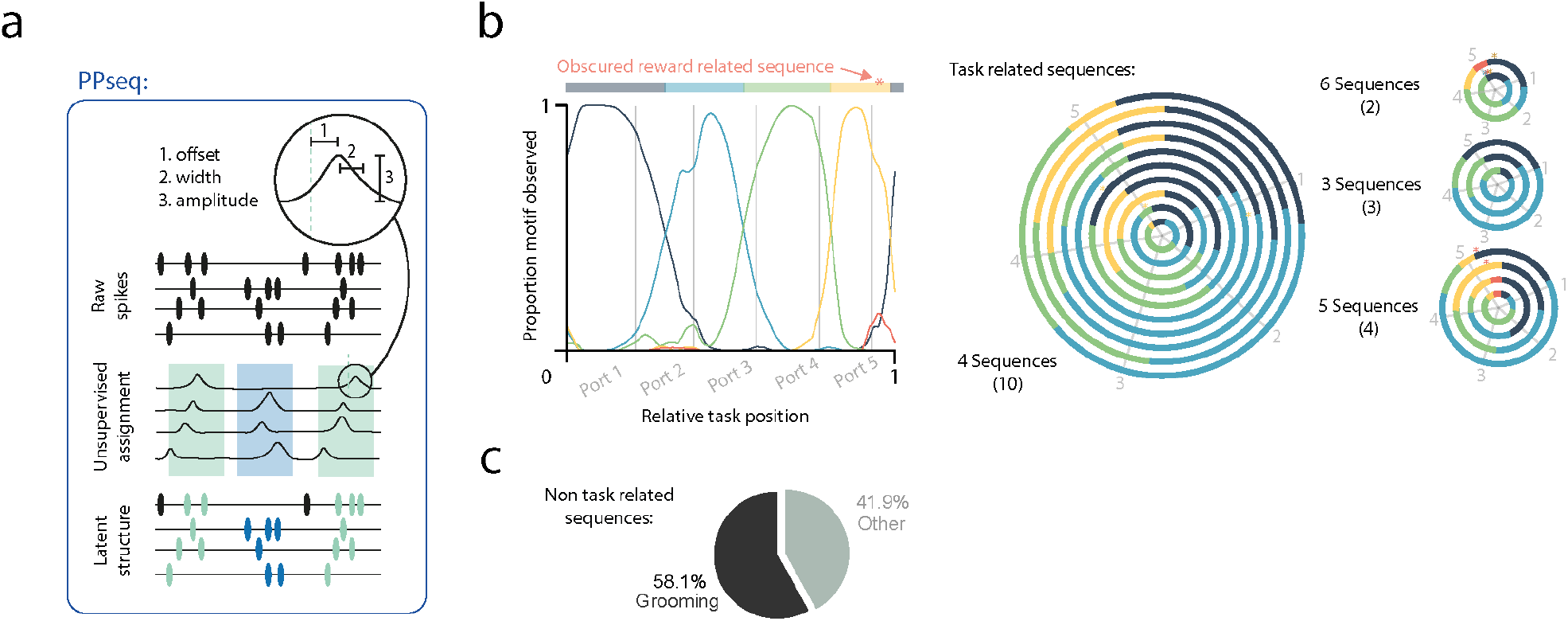
Unsupervised labelling of neural sequences during task activity. **a**. Schematic diagram outlining unsupervised detection and labelling of latent neural structure from raw spikes by PP-Seq. The model takes raw spikes, infers latent causal events which can explain repeating sequential structures within the data, then labels spikes which contribute to those events parametrised by three features of neural firing; offset, amplitude and width. **b**. Left: relative sequence incidence curves across standardised task space for an example recording session. Grey lines indicate respective task port locations across standardised space. Curves are flattened into representation of sequence incidence across task space. Colours are defined by sequences that dominate on average. A hidden (non-dominant) task relevant sequence is represented by the star. Right: same as described for left but circularised for all recordings. **c**. Percentage of non-task related sequences which were identified as related to grooming or some other feature of behaviour. (n = 8 mice, n = 19 sessions).

**Extended Data Figure 6:**
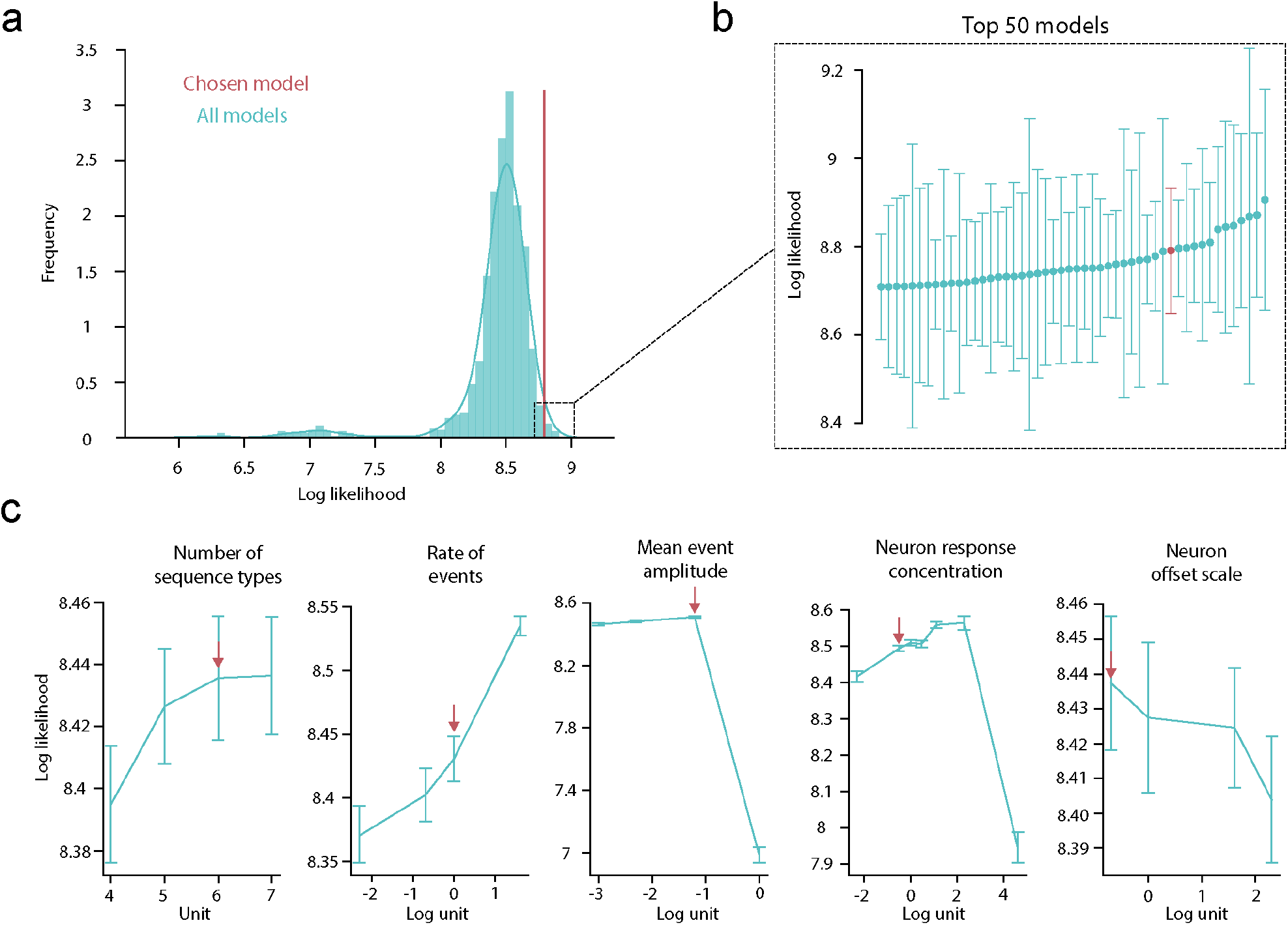
PP-Seq model hyperparameter selection. **a**. Histogram showing loss values for every combination of hyperparameters (models) tested in the grid search. The chosen model is indicated by the red line. Chosen model was hand selected from the top 20 models based on qualitative scoring of the spike labelling output **b**. Loss values and error (SEM) for the top 50 models from the search, chosen model is showing in red). **c**. Loss values as a function of hyperparameter value for each varied parameter. Error bars are SEM loss for models in which the parameter of interest was fixed and the other 5 variable hyperparameters were swept across the full range of tested values. Red arrows show the value chosen.

**Extended Data Figure 7:**
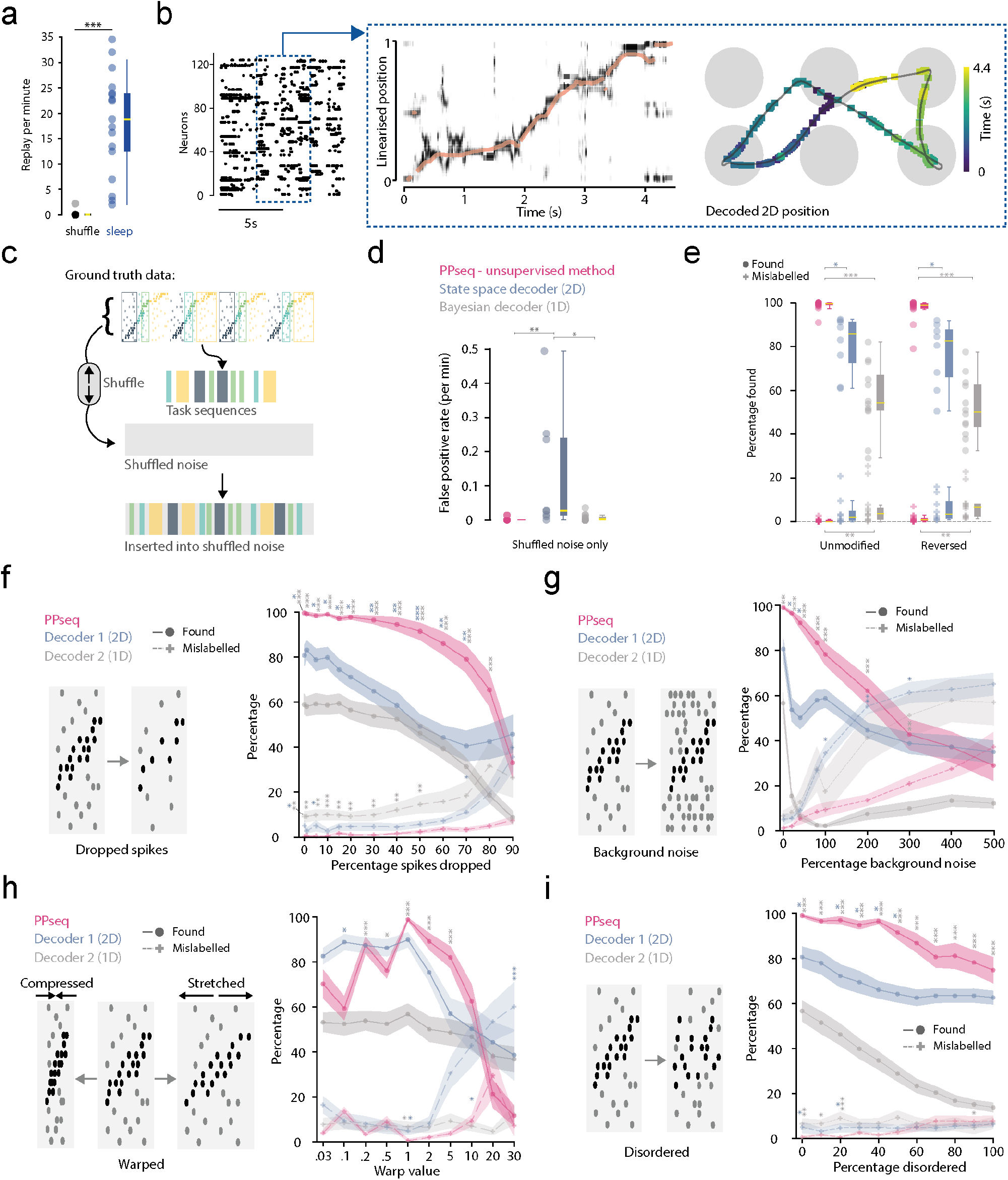
Unsupervised replay detection outperforms a state-of-the-art decoding approach on ground truth data. **a**. Average (median) replay rate per minute during sleep epochs and for shuffled data (neuron ID permuted) (For shuffle: zero false positives in 20 out 25 sessions, median rate per minute was = 0, mean = 0.18). **b**. Left: raster plot showing striatal neural activity during task execution. Blue highlighted region is an example of spiking during a single task trial. Middle: Top linearised trajectory of the task. Shading shows the posterior likelihood (white = 0, darker = more likely) across all linearised spatial bins (average in orange) Right: decoded 2D trajectory of a single trial activity, each square corresponds to the weighted average posterior probability across 2D space projected to an average trajectory of the task. **c**. Schematic showing production of ground truth ‘synthetic replay’. Time periods containing PP-seq identified neural sequences were extracted and pseudo-randomly implanted into spike matched shuffled data (neuron ID’s permuted). **d**. False positive rates (events detected per minute in background noise) for PP-Seq and the decoders (Kruskal-Wallis; p = 0.00652. post-hoc Dunn test; p = 0.0281, p = 0.0). **e**. Round markers: percentage of normal and reverse implanted sequences that were correctly identified by PP-Seq and the decoders (Unmodified, Kruskal-Wallis; p < 0.0001. Left to right, post-hoc Dunn test; p = 0.0281, p < 0.0001. Reversed, Kruskal-Wallis; p < 0.0001. Left to right, post-hoc Dunn test; p = 0.0416, p < 0.0001). Plus markers: percentage of normal and reverse implanted sequences that were identified but mislabelled as the wrong sequence type by PP-Seq and the decoders (Unmodified, Kruskal-Wallis; p = 0.00253. post-hoc Dunn test; p = 0.00224. Reversed, Kruskal-Wallis; p = 0.00141. post-hoc Dunn test; p = 0.00138). **f**. Left: schematic diagram of the type of test done: the modification done to each implanted sequence. Right, circles: for different percentage spike drop out, the percentage of sequence that were correctly identified by PP-Seq and the decoder (Kruskal-Wallis; p < 0.001, Post-hoc Dunn test; for stars (6 spines) shown left to right, p = 0.0462, p < 0.001, p < 0.001, p = 0.02971, p < 0.001, p = 0.0417, p < 0.001, p = 0.0347, p < 0.001, p = 0.0150, p < 0.001, p = 0.00476, p < 0.001, p = 0.00274, p < 0.001, p < 0.001, p < 0.001, p = 0.00114, p < 0.001, p = 0.00501, p < 0.001, p < 0.001. Right, pluses: Sequence percentages that were identified but mislabelled as the wrong sequence type by PP-Seq and the decoders (Kruskal-Wallis; p < 0.001, Post-hoc Dunn test; for stars (8 spines) shown left to right, p = 0.0247, p = 0.00242, p = 0.00403, p = 0.0258, p = 0.00108, p < 0.001, p = 0.00791, p = 0.00759, p = 0.00857, p = 0.00285, p = 0.00671, p = 0.02589, p = 0.0361). **g**. Same as d, but for percentage background noise added to sequences (Percentage found, circles: Kruskal-Wallis; p < 0.001, Post-hoc Dunn test; for stars (6 spines) shown left to right, p = 0.00414, p = 0.0160, p < 0.001, p = 0.0189, p < 0.001, p < 0.001, p < 0.001, p < 0.001, p = 0.00718. Percentage mislabelled, pluses: Kruskal-Wallis; p < 0.001, Post-hoc Dunn test; for stars (8 spines) shown left to right, p = 0.0291, p = 0.00841, p = 0.0259). **h**. Same as d, but for different sequence spike warps (Percentage found, circles: Kruskal-Wallis; p < 0.001, Post-hoc Dunn test; for stars (6 spines) shown left to right, p = 0.0403, p < 0.001, p = 0.0201, p < 0.001, p < 0.001, p < 0.001. Percentage mislabelled, pluses: Kruskal-Wallis; p < 0.001, Post-hoc Dunn test; for stars (8 spines) shown left to right, p = 0.01767, p = 0.02162, p = 0.02011, p < 0.001). **i**. Same as d, but for percentage disordered spikes (Percentage found, circles: Kruskal-Wallis; p < 0.001, Post-hoc Dunn test; for stars (6 spines) shown left to right, p = 0.0406, p < 0.001, p < 0.001, p = 0.0171, p < 0.001, p = 0.0256, p < 0.001, p = 0.0102, p < 0.001, p = 0.0214, p < 0.001, p < 0.001, p < 0.001, p < 0.001, p < 0.001, p < 0.001. Percentage mislabelled, pluses: Kruskal-Wallis; p < 0.001, Post-hoc Dunn test; for stars (8 spines) shown left to right, p = 0.0235, p = 0.00218, p = 0.0173, p = 0.0237, p < 0.001, p = 0.0271). For plots c-h, for each test value and for each group (PP-Seq and Decoders), n = 10 sessions from n = 6 implanted animals.

**Extended Data Figure 8:**
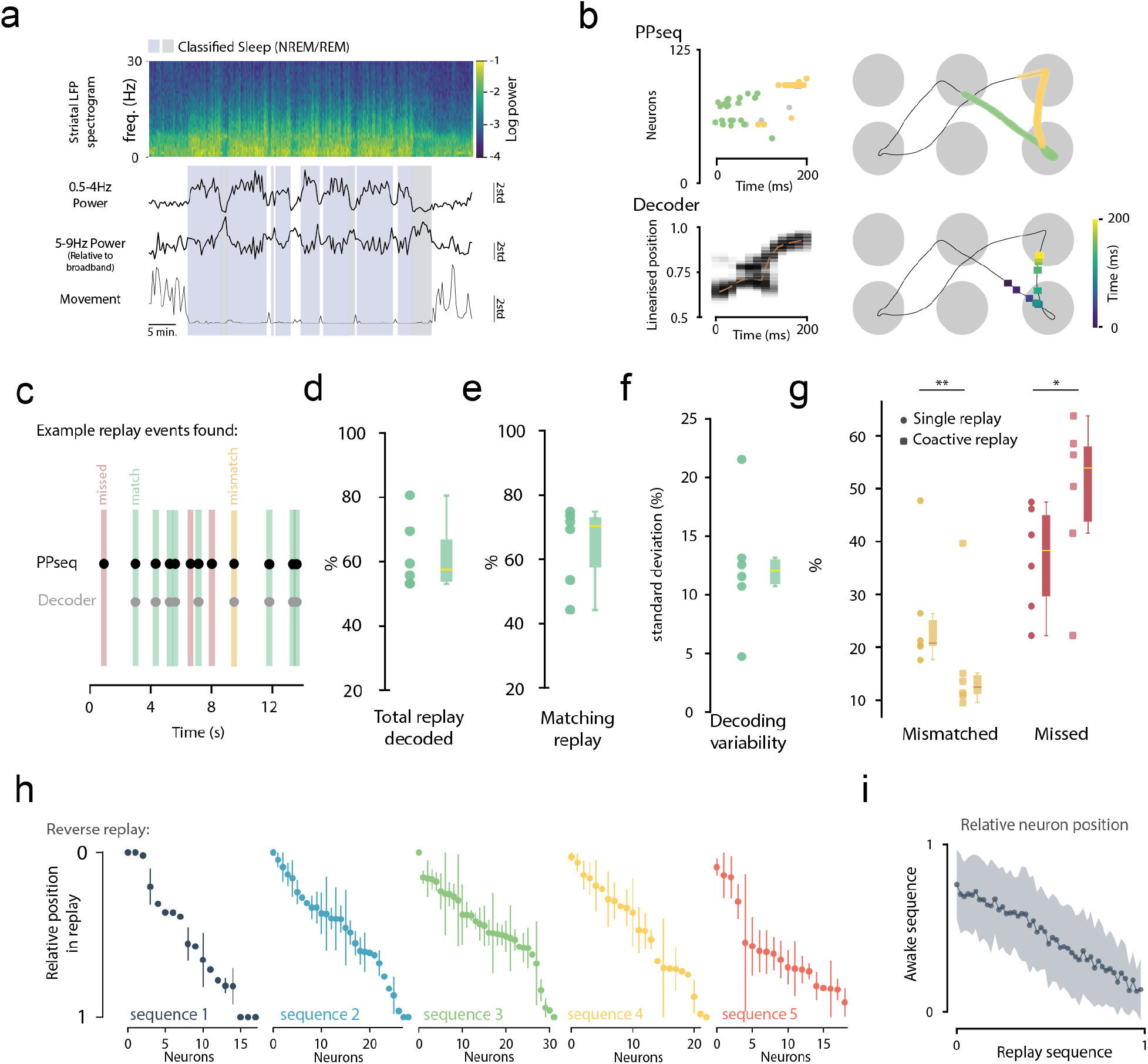
Characteristics and decodability of procedural replay in the striatum. **a**. Spectrogram for an example sleep epoch with Delta and Theta band spectral power and average video tracking movement velocity underneath (traces). Sleep periods were NMREM or REM periods classified from these metrics (see methods). **b**. Top: PP-Seq labelled spikes (left) and corresponding PP-seq sequence positions observed during task activity projected onto average tracking trajectory (right). Bottom: (left) for the spikes shown above, the decoded 1D position across linearised task space (red dashed line). Shading shows the posterior likelihood across all spatial bins (white = 0, darker = more likely). Maximum of the 1D decoded position (the most probable position) projected onto 2D average tracking trajectory (right). **c**. Example epoch showing PP-Seq identified replay events and replays that were also found by the decoder. Perfect matches (green markers), replays with mismatched spatial locations (yellow) and PP-seq replay not found by the decoder (red) are shown. **d**. Percentage of PP-Seq replay events that were also found by the decoder. **e**. The percentage of found events which were spatially harmonious between the two methods. **f**. Standard deviation of total found percentages between sequence types for each session. **g**. Percentage mismatched and missed sequences for decoded epochs which contained either a single PP-Seq sequence (circle) or multiple coactive sequences (square) (n = 6 sessions, n = 6 mice. paired t-test; p = 0.001, p = 0.01). **h**. Mean relative position of spikes, for neurons in each reversed replay sequence observed in an example recording (error bars, SEM). **i**. Mean relative positions in awake sequences vs. reverse replay sequences for all analysed neurons across every session (OLS regression, r = -0.58, p < 0.001).

**Extended Data Figure 9:**
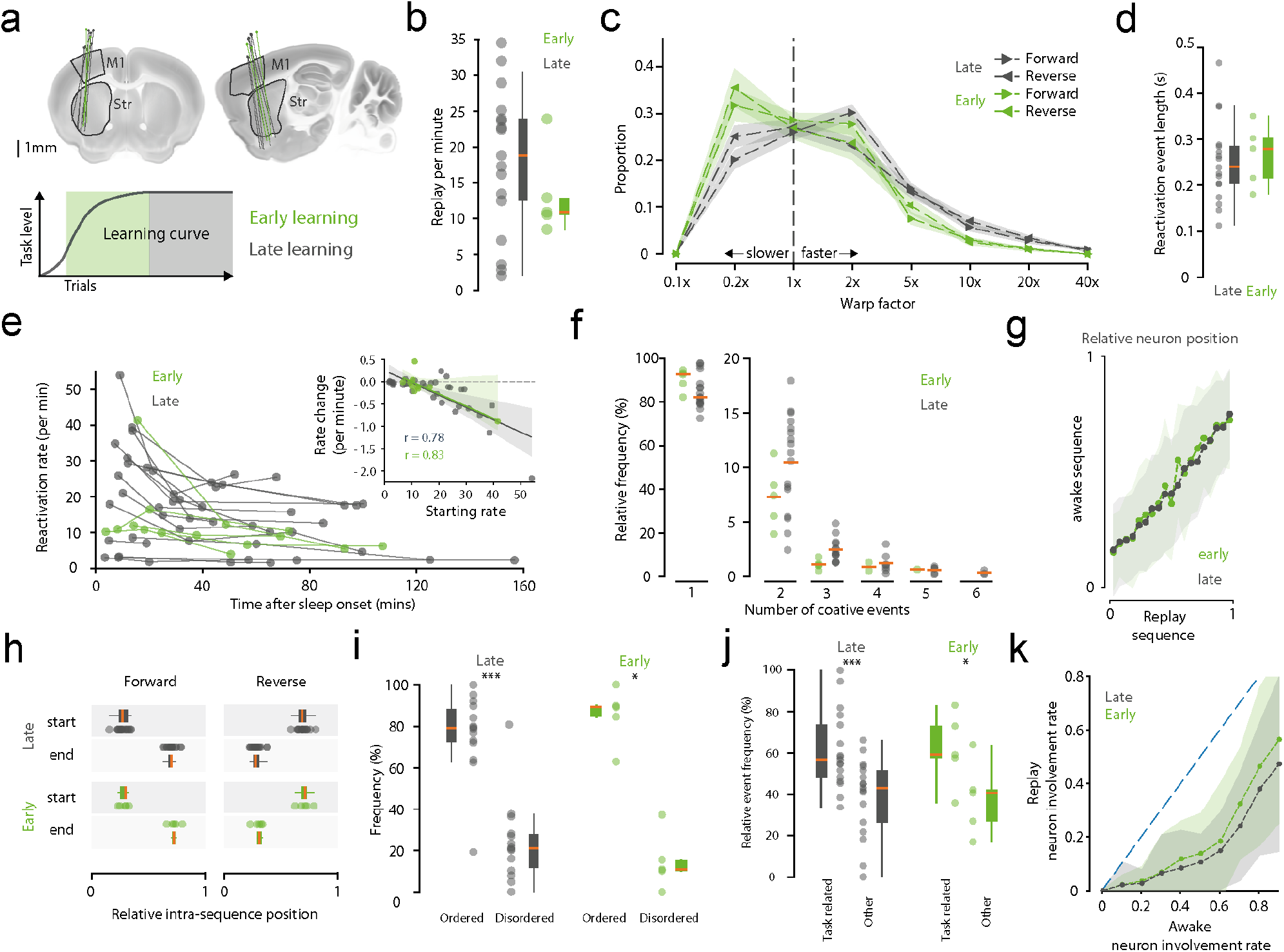
Procedural replay is independent of learning stage. **a**. Top: schematic showing implanted neuropixel probe locations projected onto standard Allen atlas coronal (left) and sagittal (right). Bottom: schematic diagram showing early learning (before level 50) and late learning (after level 50) recording groups. **b**. Replay event rate per minute (independent t-test, p = 0.235). **c**. Distribution of warp factors for forwards and backwards replay events (1x represents awake speed).(Differences for all warp factors, Kruskal-Wallis; p <0.0001. Paired difference for early and late, forward warp factor groups left to right, post-hoc Dunn test; p = 1.0, p = 0.216, p = 0.938, p = 0.724, p = 0.323, p = 0.684, p = 0.830, p = 0.596. Paired differences for early and late, reversed warp factor groups left to right, post-hoc Dunn test; p = 1.0, p = 0.271, p = 0.818, p = 0.997, p = 0.770, p = 0.590, p = 0.177, p = 0.689). **d**. Average (mean) single replay event lengths (duration from first to last spike) for each recording group (independent t-test, p = 0.995). **e**. Main: reactivation rates for each analysed sleep epoch against time from first sleep onset. (Multivariate comparison between groups, MANOVA; Wilks lambda, p = 0.961) Inset: rate change against starting rate for each pair of epochs per session. **f**. Frequency of single (isolated) and coactive events for each recording session (Differences between all groups, Kruskal-Wallis; p<0.001. Paired differences between early and late for each group, left to right, post-hoc Dunn test; p = 0.571, p =0.650, p=0.299, p=0.896, p=0.961). **g**. Mean relative positions in awake sequences vs. reverse replay sequences for all analysed neurons across every session (Multivariate comparison between groups, MANOVA; Wilks lambda, p = 0.9756). **h**. Average (mean) start and end points for all forward (top) and reverse (bottom) replay events. Position is relative to the corresponding average awake sequence. (Comparison between all groups, Kruskal-Wallis; p <0.001. Pairwise comparison between forward start groups, post-hoc Dunn test; p = 0.876. Pairwise comparison between forward end groups, post-hoc Dunn test; p = 0.596. Pairwise comparison between reverse start groups, post-hoc Dunn test; p = 0.806). Pairwise comparison between reverse end groups, post-hoc Dunn test; p = 0.880). **i**. Relative frequencies of task ordered and disordered coactive sequences (Early, ordered difference from 62.04%: chance level, Wilcoxon signed-rank; p =0.0655. Late, ordered difference from 62.04%: chance level, Wilcoxon signed-rank; p =0.0215. Early, difference between subgroups, permutation test: observed difference in means = 70.5%, 99^th^ percentile permuted difference = 51.6%, p < 0.00386. Late, difference between subgroups, permutation test: observed difference in means = 50.3%, 99^th^ percentile permuted difference = 29.2%, p < 0.001. Pairwise difference for Early and Late ordered sequences, MannWhitney U test; p = 0.259).**j**. Normalised percentages of task and non-task related replay observed (Pairwise difference for early and late task related sequences, MannWhitney U test; p = 0.317). (Early, task related difference from chance level: 50%, one sample t-test; p = 0.103. Late, task related difference from chance level: 50%, one sample t-test; p < 0.0001. Early, difference between subgroups, Permutation test: observed difference in means = 36.4%, 99^th^ percentile permuted difference = 27.2%, p = 0.0113. Late, difference between subgroups, Permutation test: observed difference in means = 21.20%, 99^th^ percentile permuted difference = 15.88%, p = 0.0004. Pairwise difference for Early and Late task related sequences; independent t-test, p = 0.317). **k**. Relative individual neuron involvement frequencies for each sequence during awake task activity and sleep periods (Multivariate comparison between groups, MANOVA; Wilks lambda, p = 0.704). (Late: n = 19 sessions, n = 8 mice, Early: n = 5 sessions, n = 3 mice).

**Extended Data Figure 10:**
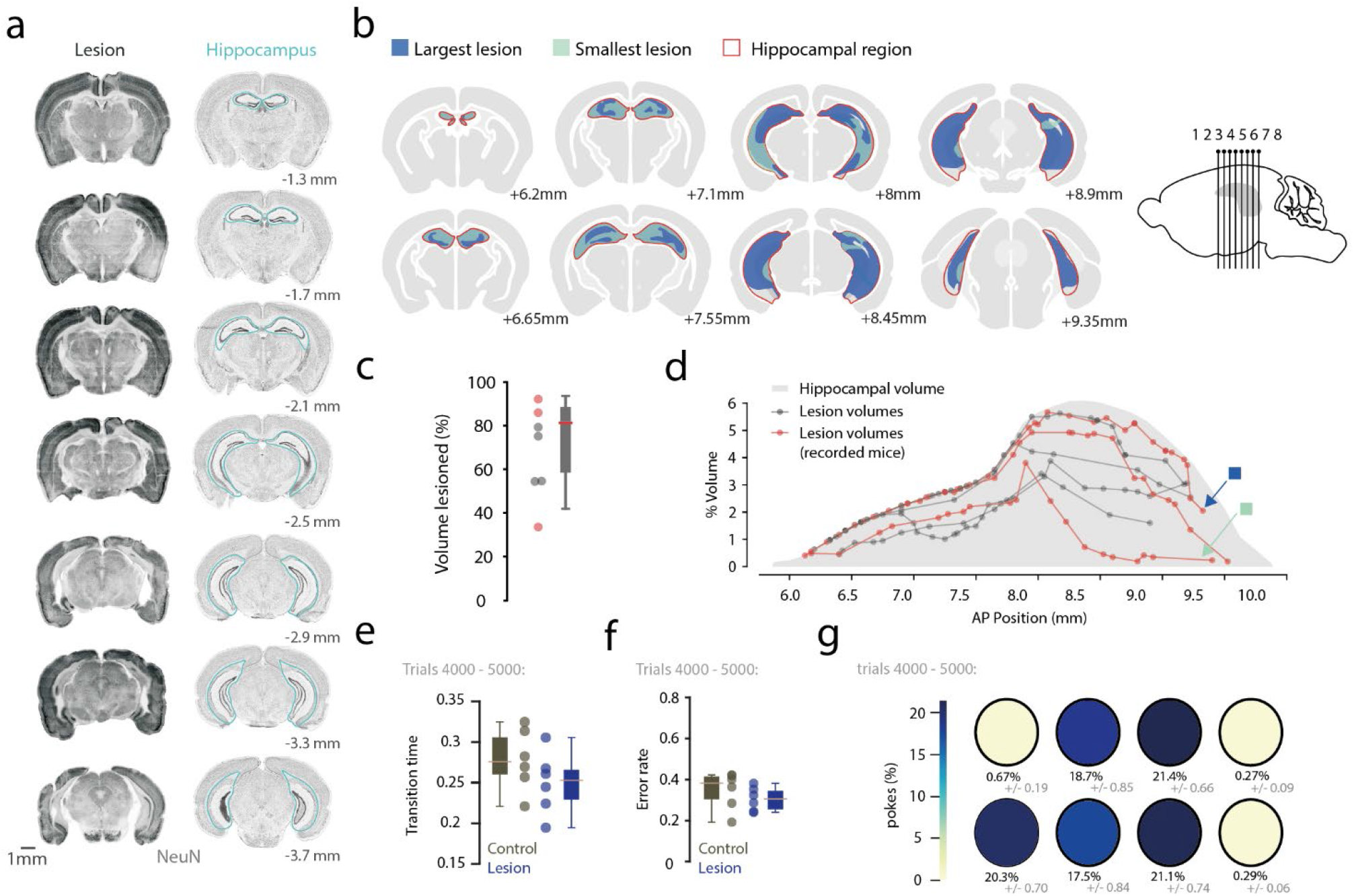
Hippocampus lesion extent. **a**. Left: histology for example NeuN stained coronal slices showing lesion extent across hippocampal volume. Right: corresponding Allen reference atlas slices with hippocampal volume highlighted (blue outline). **b**. Largest (blue shaded) and smallest (green shaded) lesion extent for coronal slices across the extent of the hippocampus. **c**. Percentage of total hippocampal volume lesioned for each animal. Mice that were used in neuropixel recording experiments are shown in red. **d**. Percentage of total hippocampal volume lesioned for each animal across the extent of the hippocampus. Total hippocampal volume is shown (grey). Smallest and largest lesions are labelled by blue and green markers respectively. Mice that were used in neuropixel recording experiments are shown in red (n = 7 mice). **e**. Port-to-port transition errors for expert (trials 4000-5000) mice in lesion and control groups. (Independent t-test, p = 0.2251) **f**. Average (mean) transition port-to-port transition times for expert mice in each group. (Wilcoxon rank sum, p = 0.3597) **g**. Average (mean) percentage port poke occurrences for mice in the lesion group (SEM shown in grey, control n = 6 mice, lesion n = 6 mice).

**Extended Data Figure 11:**
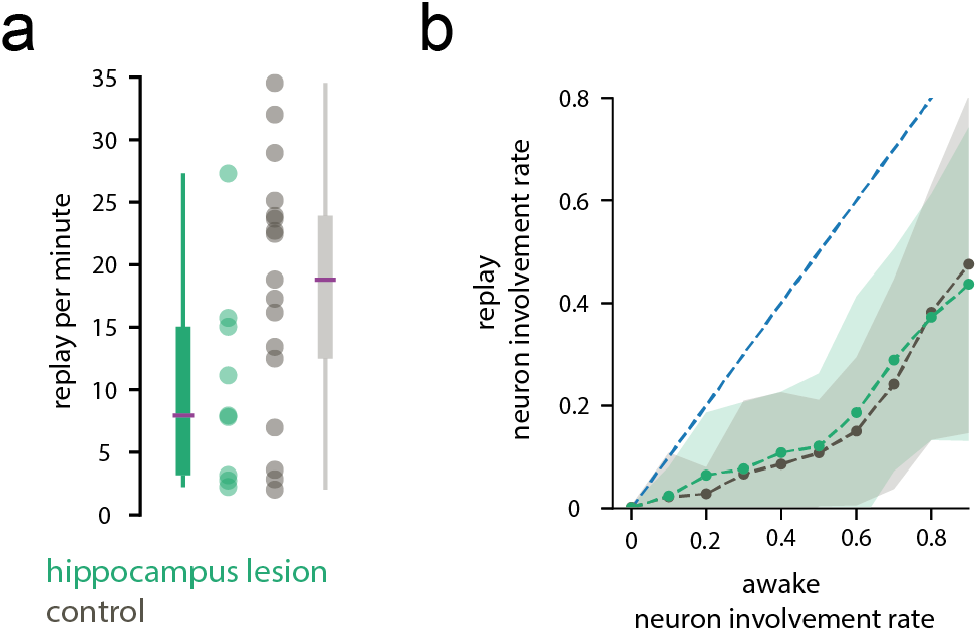
Procedural replay characteristics are unchanged by bilateral lesion to the hippocampus. **a**. Replay event rate per minute for lesion and control time epochs (independent t-test p =0.109). **b**. Relative individual neuron involvement frequencies for each sequence during awake task activity and sleep periods (Multivariate comparison between groups, MANOVA; Wilks lambda, p = 0.938). (Control: n = 19 sessions, n = 8 mice, lesion: n = 9 sessions, n = 3 mice).

## Notes

### Competing Interest Statement

The authors have declared no competing interest.

